# Chromatin structure of the inactive X chromosome revealed by in situ cryo-electron tomography

**DOI:** 10.64898/2026.09.07.749863

**Authors:** Jeongyoon Choi, Dimitrios Bellos, Joseph Bowness, Adam Cawte, Neil Brockdorff

## Abstract

The intricate organisation of chromatin in eukaryotic cells plays a central role in regulating gene transcription and other DNA-templated processes. Prior studies using light microscopy, biochemistry, and genomics have implicated histone and DNA modifications, histone variants, chromosomal proteins, non-coding RNA, and intrinsic biophysical properties of chromatin in determining chromatin organisation across multiple scales. In this study, we apply an innovative approach, direct visualisation of native chromatin structure and its modulation by different factors using cryo-electron tomography, exploiting the inactive X chromosome (Xi) in differentiating mouse embryonic stem cells as a model system of facultative heterochromatin. We show that Xi chromatin undergoes progressive compaction across scales, from individual nucleosomes to chromatin domains. We observe an enrichment of histone H1-bound chromatosomes that are distributed to form the core of Xi chromatin domains. Moreover, we demonstrate that histone deacetylation demarcates chromatin domains. Specifically, perturbations leading to histone acetylation in the Xi dissolve discrete chromatin-domain boundaries, resulting in a homogeneous chromatin distribution with a dense and uniform packing of nucleosomes. Together, these findings illuminate the multiscale organisation of native facultative heterochromatin and the contributions of histone H1 and histone acetylation to this organisation. This study opens a new avenue for investigating how chromatin modification and structure relate to gene activity in near-native cells.

---

DNA in eukaryotic cells is organised in a hierarchical manner, ranging from the scale of the nucleosome (∼10 nm), nanoscale chromatin domains (50–200 nm), topologically associated domains (TADs, ∼1 µm), to chromosome territories (a few µm)^1^. These different levels of chromatin organisation play a critical role in regulating DNA-templated processes, including gene transcription. This concept was first proposed in the context of classical light microscopy studies that defined distinct open and compact chromatin states, euchromatin and heterochromatin, subsequently shown to correspond to gene-rich versus transcriptionally silent genomic regions^2^. A further distinction that emerged from early studies is between constitutive and facultative heterochromatin, the latter referring to regions of the genome that switch between euchromatic and heterochromatic states in response to specific signals, for example, during development. A classical example of facultative heterochromatin is the inactive X chromosome (Xi) in female mammalian cells. Xi is established by X-chromosome inactivation (XCI), a stepwise process that initiates in cells of early embryos and is orchestrated by the non-coding RNA Xist. Specifically, Xist RNA engages factors that trigger a cascade of biochemical and structural changes in chromatin, modifying chromatin organisation at both the nucleosome scale and higher-order levels, and leading ultimately to Xi chromatin compaction and chromosome-wide gene silencing^3–5^.

Key advances towards understanding chromatin organisation and how it relates to gene activity have come from cellular imaging, biochemical, and genomics-based assays^6^. A recently developed methodology is in situ structural biology, in which the native structure of chromatin is determined using cryo-electron tomography (cryo-ET)^7–9^. This approach offers critical advantages in that it provides resolution across scales from individual nucleosomes to sub-chromosome territories, whilst conserving near-native structure and organisation. In this study, we expand the application of cryo-ET to investigate chromatin structural changes associated with distinct transcriptional states, using the establishment of XCI as a model system. Thus, we apply cryo-correlative light and electron microscopy (cryo-CLEM) combined with targeted cryo-focused ion beam (FIB) milling of the Xi chromosome territory to investigate both the stepwise maturation of Xi in differentiating mouse embryonic stem cells (mESCs) and the effect of perturbing histone deacetylation, a key Xi silencing pathway.

## In situ chromatin structure of the native Xi

To investigate the native chromatin structure of Xi, we established a targeted cryo-ET analysis in differentiating female embryo-derived (XX) mESCs, using cryo-CLEM combined with cryo-FIB milling (Extended Data Fig. 1A and see Methods). Thus, we modified an existing interspecific XX mESC line with doxycycline-inducible Xist expression^10^, introducing an mCherry-fused Ciz1 transgene^11^, to enable identification of the Xi using cryo-light microscopy (cryo-LM) (Extended Data Fig. 1A, B). We confirmed Xist-mediated X-linked gene silencing using chromatin-associated RNA sequencing (ChrRNA-seq) analysis^10^ (Extended Data Fig. 1C).

We went on to acquire cryo-ET tilt-series data centred on Xi territories from day-7 differentiated cells, at which time Xi gene silencing is mostly complete (Extended Data Fig. 1C). A representative Xi tomogram is shown in Fig. 1A and Supplementary Video 1. We observed nucleosomes forming clusters at scales comparable to nanoscale chromatin domains (50–200 nm)^1^ and separated by chromatin-depleted regions, or interchromatin space (Fig. 1A, Fig. 1B left panel, and Supplementary Video 1). We were also able to discern heterogeneous non-chromatin densities that exhibit a slightly lower contrast than that of nucleosomes (Fig. 1A, C). We speculate that these densities likely correspond to ribonucleoprotein (RNP) particles (see Discussion).

**Figure 1.**
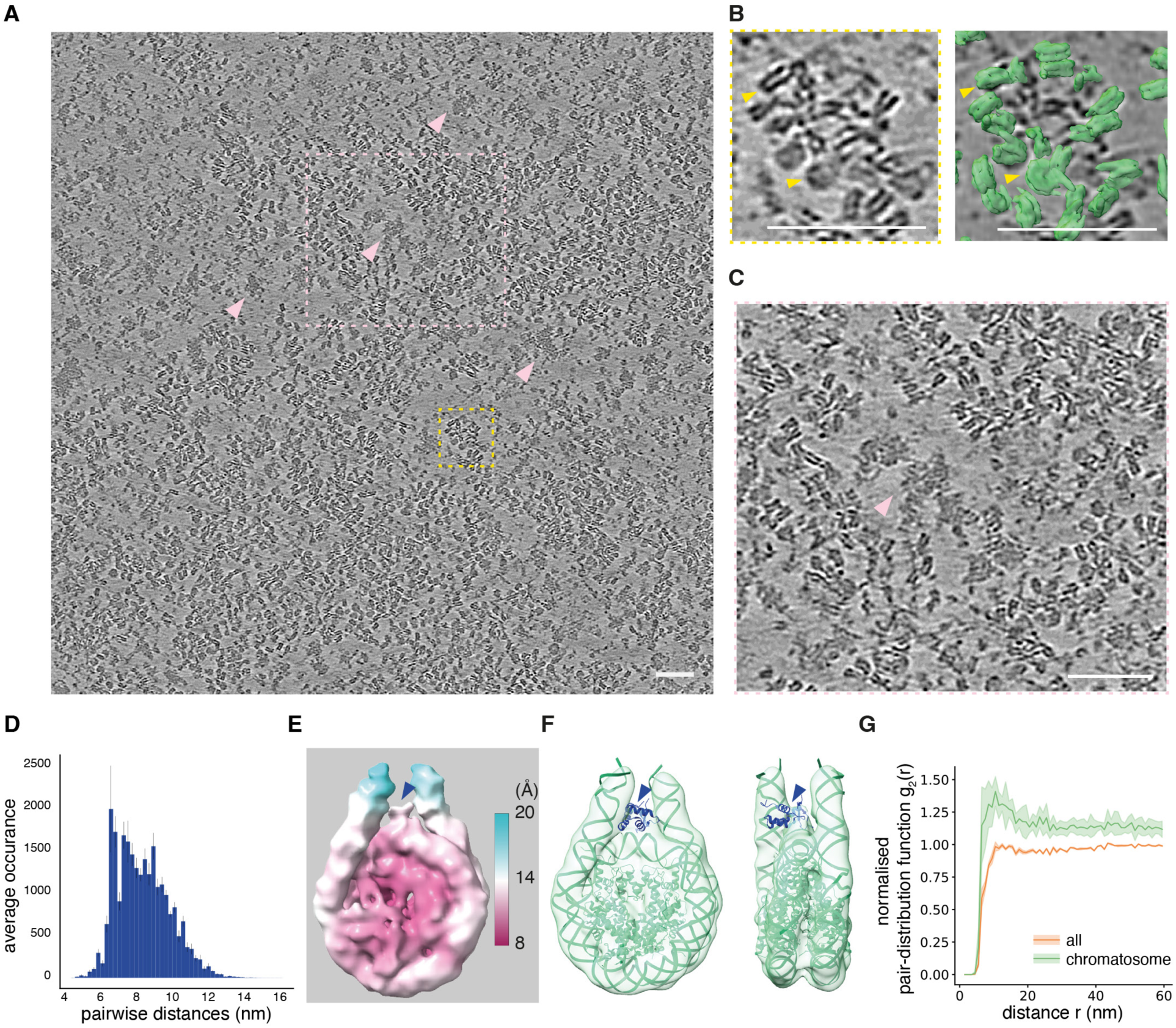
In situ chromatin structure of the inactive X chromosome in differentiated XX mESCs. **(A)** An Xi tomogram slice from a day-7 differentiated XX mESC; scale bar = 50 nm. Pink arrows indicate potential RNPs residing in chromatin-depleted regions. **(B)** Left: A zoomed view of the yellow-boxed area in (**A**); yellow arrows indicate nucleosomes in side view and top view; scale bar = 50 nm. Right: the same zoomed area with an overlay of extracted nucleosome particles in green, picked by optimised template matching. **(C)** A zoomed view of the pink-boxed area in (**A**); pink arrows indicate putative RNPs; scale bar = 50 nm. **(D)** A histogram of averaged pairwise distances between NN nucleosomes of the Xi across 5 tomograms. Error bars represent standard deviation (SD). **(E)** STA map of the chromatosome derived from Xi tomograms, coloured by local resolution (key on the right). The blue arrow indicates putative histone H1 density. **(F)** The human chromatosome model (PDB: 7DBP) was fitted into our STA map. Extra cryo-EM density corresponding to histone H1 (shown in blue in the fitted model) is indicated by arrows. **(G)** A plot of the averaged, normalised pair-distribution function g_2_(r) of all nucleosomes and chromatosomes, up to a distance r of 60 nm, in increments of 1 nm. A solid line represents the mean, and shading represents the SD, across the 5 Xi tomograms.

To analyse chromatin organisation quantitatively, we optimised nucleosome particle picking by employing a local thresholding algorithm for particle extraction, and by combining particles picked from two independent template-matching runs performed on raw and missing-wedge-predicted tomograms (Extended Data Fig. 2A and Fig. 1B right panel). The histogram of pairwise distances between nearest-neighbour (NN) nucleosomes (Fig. 1D) showed a distribution comparable to that reported in a cryo-tomogram analysis of human chromatin, in which every nucleosome was manually segmented^12^. Mean volume fraction occupied by nucleosomes (Extended Data Fig. 2B) fell within a range comparable to that reported in a quantitative fluorescence and electron spectroscopic imaging study^13^, further supporting the reliability of our optimised nucleosome particle picking. Superimposition of tomograms with the location of extracted nucleosome particles highlights chromatin depletion in the regions surrounding putative Xi RNPs (Extended Data Fig. 2C).

By performing 3D classification and refinement of the extracted nucleosome particles from five Xi tomograms, we obtained a subtomogram averaging (STA) map of Xi nucleosomes (Fig. 1E and Extended Data Table 1). Notably, the nucleosome class with sufficiently homogeneous structural conformation to yield an STA map exhibited clear histone H1 density, forming a chromatosome (Fig. 1F). Interestingly, in pairwise angular orientation analysis of NN nucleosomes, the nucleosome particles that yielded the chromatosome STA map exhibited a significantly lower proportion of face-to-side arrangement than that of all nucleosomes (Extended Data Fig. 3A, B), implying that H1-bound chromatosomes are more likely to adopt a stacked or ’zig-zag’-like arrangement. We also found that chromatosomes form clusters in 3D when mapped back to the tomogram (Extended Data Fig. 3C), showing clear overlap with the core of nanoscale chromatin domains (Extended Data Fig. 3D). That chromatosomes tend to co-cluster is further demonstrated by the normalised pair-distribution function g_2_(r) being above 1 at distances between 7 nm and 15 nm (Fig. 1G). In contrast, g_2_(r) for all extracted nucleosome particles had a value of ∼1 for all distances.

## Chromatin compaction of the native Xi resolved at sub-diffraction scales during XCI progression

We went on to apply our cryo-CLEM pipeline to quantify the features of Xi chromatin during the establishment of XCI in differentiating XX mESCs. Prior studies have revealed that Xi chromatin is established through distinct stages, with histone deacetylation and deposition of Polycomb-mediated histone modifications occurring in step with the onset of Xist RNA expression, and with structural changes at the TAD scale, global chromatin compaction, and features such as CpG island DNA methylation, emerging several days later^5^. The chromosomal protein SMCHD1 plays a central role in instigating late-stage modification of Xi chromatin^14^. SMCHD1 enrichment over Xi occurs progressively, peaking between day-5 and day-7 of differentiation^11,15^. Accordingly, we set out to examine native Xi chromatin reorganisation during XCI progression, quantifying features at the chromatin-domain level and below, corresponding to sub-diffraction-limit scales (below 200 nm) both at day-5 and day-7 of differentiation. As a control, we reconstructed tomograms from randomly chosen nuclear regions lacking mCherry-CIZ1 fluorescent signal, referred to henceforth as other chromosomes. Quantitative chromatin feature and STA analyses were performed as described above using 26 tomograms.

Representative tomograms from day-5 and day-7 differentiated mESCs are shown in Extended Data Fig. 4A, B, Extended Data Fig. 5A, B and Supplementary Video 1-4. In these examples, other chromosome regions display more dispersed chromatin domains, with larger chromatin-depleted regions, compared with matched Xi tomograms, consistent with the prediction that they include a random distribution of euchromatin and heterochromatin domains. Putative RNPs were observed in both Xi and other chromosome tomograms but were more numerous and spatially separated from chromatin domains in the latter. As noted for day-7 Xi nucleosomes, the STA map of day-5 Xi nucleosomes contained extra density corresponding to histone H1 (Extended Data Fig. 5C), indicating that histone H1 enrichment is an early-stage Xi modification. H1 density was not evident in the STA map of nucleosomes from other chromosome tomograms (Extended Data Fig. 5D).

Quantification of chromatin features at the nucleosomal scale revealed that mean volume fraction occupied by nucleosomes increased ∼1.4-fold in the Xi of day-7 cells compared with the Xi of day-5 cells (Fig. 2A), similar to a previous report of ∼1.2-fold global compaction of the Xi measured by DNA-FISH in human cells^16^. In addition, the day-7 Xi exhibited higher coordination number^17^ (Fig. 2B), defined here as the number of neighbouring nucleosomes per nucleosome within a sphere of 12 nm radius. We also observed that the day-7 Xi exhibited shorter pairwise distances between NN nucleosomes (Fig. 2C) and a smaller effective volume occupied by each nucleosome, indicated by higher Voronoi-based local density^18,19^ (Fig. 2D). Together, these results demonstrate that the day-7 Xi chromatin undergoes moderate but statistically significant compaction at the single-nucleosome level during XCI progression.

**Figure 2.**
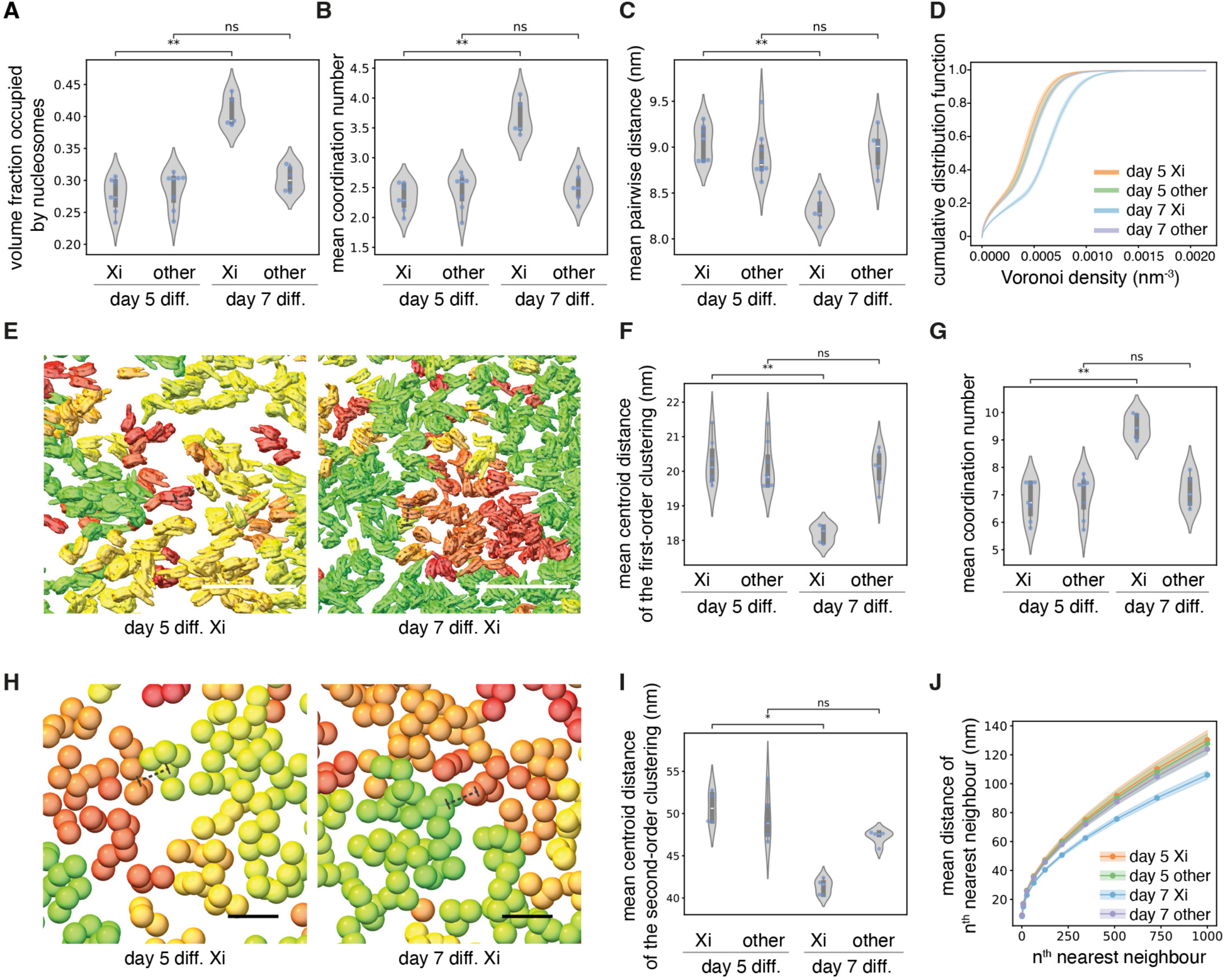
The inactive X chromosome undergoes moderate chromatin compaction across scales during XCI progression. Chromatin features determined for day-5 and day-7 differentiated XX mESCs: Violin plots of **(A)** volume fraction occupied by nucleosomes, **(B)** mean coordination number of nucleosomes within a sphere of 12 nm radius, and **(C)** mean pairwise distance between NN nucleosomes. **(D)** A plot of cumulative distribution function (CDF) of Voronoi-based local density. The line represents the mean, and the shaded region represents the standard error of the mean across tomograms. **(E)** Representative cropped views showing nucleosome clustering of the day-5 Xi (left) and day-7 Xi (right) within the 3D tomogram volume (z = 46.6 nm). Nucleosomes belonging to the same cluster are depicted in the same colour shade. Violin plots of **(F)** mean pairwise distance between centroids of NN first-order clusters, as indicated by dotted lines in **(E)**; and **(G)** mean coordination number of nucleosome clutches within a sphere of 36 nm radius, based on the centroids of the first-order clusters. **(H)** Representative cropped views showing second-order clustering of first-order clusters within the 3D tomogram volume (z = 46.6 nm). Each first-order cluster is depicted as a sphere with a diameter of 28 nm around the centroid of the cluster, for simplification. First-order clusters belonging to the same second-order cluster are depicted in the same colour shade. **(I)** Mean pairwise distance between centroids of NN second-order clusters, as indicated by dotted lines in (**H**). **(J)** Mean pairwise distance between n^th^ NN nucleosomes. Solid lines represent the mean, and shading represents the SD. Scale bar = 50 nm for (**E**) and (**H**). Each blue dot in plots (**A**)–(**C**), (**F**), (**G**), and (**I**) represents a data point from a subvolume (839 × 839 × 46.6 nm^3^) of an independent tomogram, consisting of >20,000 extracted nucleosome particles. (**D**) and (**J**) use the same n. n = 7 (day-5, Xi); 8 (day-5, other); 5 (day-7, Xi); 6 (day-7, other). Exact P-values between all data points are provided in Supplementary Data Table 1.

Next, to examine chromatin structural changes in the Xi at the chromatin-domain scale during XCI, we performed nucleosome clustering analysis using the hierarchical density-based spatial clustering of applications with noise (H-DBSCAN) algorithm^20^. We found that nucleosomes formed clusters (Fig. 2E, Extended Data Fig. 6A, B) with an average of ∼7 nucleosomes per cluster (Extended Data Fig. 6C). This scale is comparable to the smallest nucleosome cluster unit reported to date, previously termed nucleosome clutches^21^. Notably, mean pairwise distances of clustered NN nucleosomes (Extended Data Fig. 6D) were slightly shorter than those of all NN nucleosomes (including non-clustered ones) (Fig. 2C), with the day-7 Xi again showing the shortest mean pairwise distances among all four chromosome categories. At the nucleosome-clutch scale, the day-7 Xi exhibited shorter spacing between NN clutches (Fig. 2F) and higher mean coordination number (defined as the number of neighbouring clutches per clutch within a sphere of 36 nm radius) than the day-5 Xi (Fig. 2G). In addition, the day-7 Xi contained more clutches per unit tomogram volume than the day-5 Xi (Extended Data Fig. 6E).

We then applied the H-DBSCAN algorithm to the nucleosome clutches to investigate chromatin organisation at the second-order clustering level (Fig. 2H, Extended Data Fig. 6F, G). Each second-order cluster contained an average of ∼8 first-order clusters (i.e., nucleosome clutches) (Extended Data Fig. 6H), totalling ∼56 nucleosomes, corresponding in scale towards the lower end of the nanoscale chromatin domain range^1^. At the nanoscale chromatin-domain level, the day-7 Xi chromatin again showed shorter spacing between NN clusters (Fig. 2I) and a higher number of clusters per unit tomogram volume (Extended Data Fig. 6I). Moreover, mean pairwise distances between the n^th^ NN nucleosomes were shorter up to the 1000^th^ NN nucleosomes (Fig. 2J), and mean coordination number within spheres of increasing radius up to 132 nm was higher (Extended Data Fig. 6J), in the day-7 Xi than in the day-5 Xi.

Collectively, these results demonstrate that the native Xi chromatin undergoes progressive compaction across scales, from single nucleosomes to chromatin domains, during the establishment of the fully silenced chromosome.

## Histone deacetylation demarcates chromatin domains in the Xi

A key feature of Xi is the deacetylation of multiple lysines on histone tails^22–24^. Xi histone deacetylation is driven by the Xist RNA-interacting protein SPEN, which interacts with the NCoR/SMRT corepressor to activate histone deacetylase 3 (HDAC3)^25,26^. HDAC3 is pre-bound to putative enhancer regions that undergo histone deacetylation during XCI^27^. In prior work, we used CRISPR-Cas9 mutagenesis in interspecific XX mESCs to introduce endogenous point mutations (R3552A and R3554A) in the SPEN SPOC domain (Spen^SPOCmut^)^15^, disrupting the interaction with NCoR/SMRT^28^, and demonstrated that this results in a significant deficit in Xi gene silencing^15^. Consistent with this finding, chromatin immunoprecipitation sequencing (ChIP-seq) analysis of SPEN^SPOCmut^ demonstrates that levels of acetylated histone H3 lysine 27 (H3K27ac) at Xi gene enhancers and promoters are equivalent to those at the corresponding active X chromosome (Xa) allele (Fig. 3A, Extended Data Fig. 7A). Thus, to investigate the effect of histone deacetylation on native chromatin structure of the Xi, we re-engineered the R3552A and R3554A mutations into the SPEN SPOC domain in the XX mESC line used above for cryo-CLEM. We confirmed that the Spen^SPOCmut^ mESCs were impaired in Xi gene silencing by chrRNA-seq analysis (Fig. 3B) and in histone deacetylation using immunofluorescence (IF) analysis to determine H3K27ac, H4K16ac, and H3K9/K14ac levels within Xi territories at interphase (Fig. 3C, Extended Data Fig. 7B, C). IF analysis additionally demonstrated retention of Polycomb-mediated histone modification on the Xi in Spen^SPOCmut^ mESCs (Extended Data Fig. 7D).

**Figure 3.**
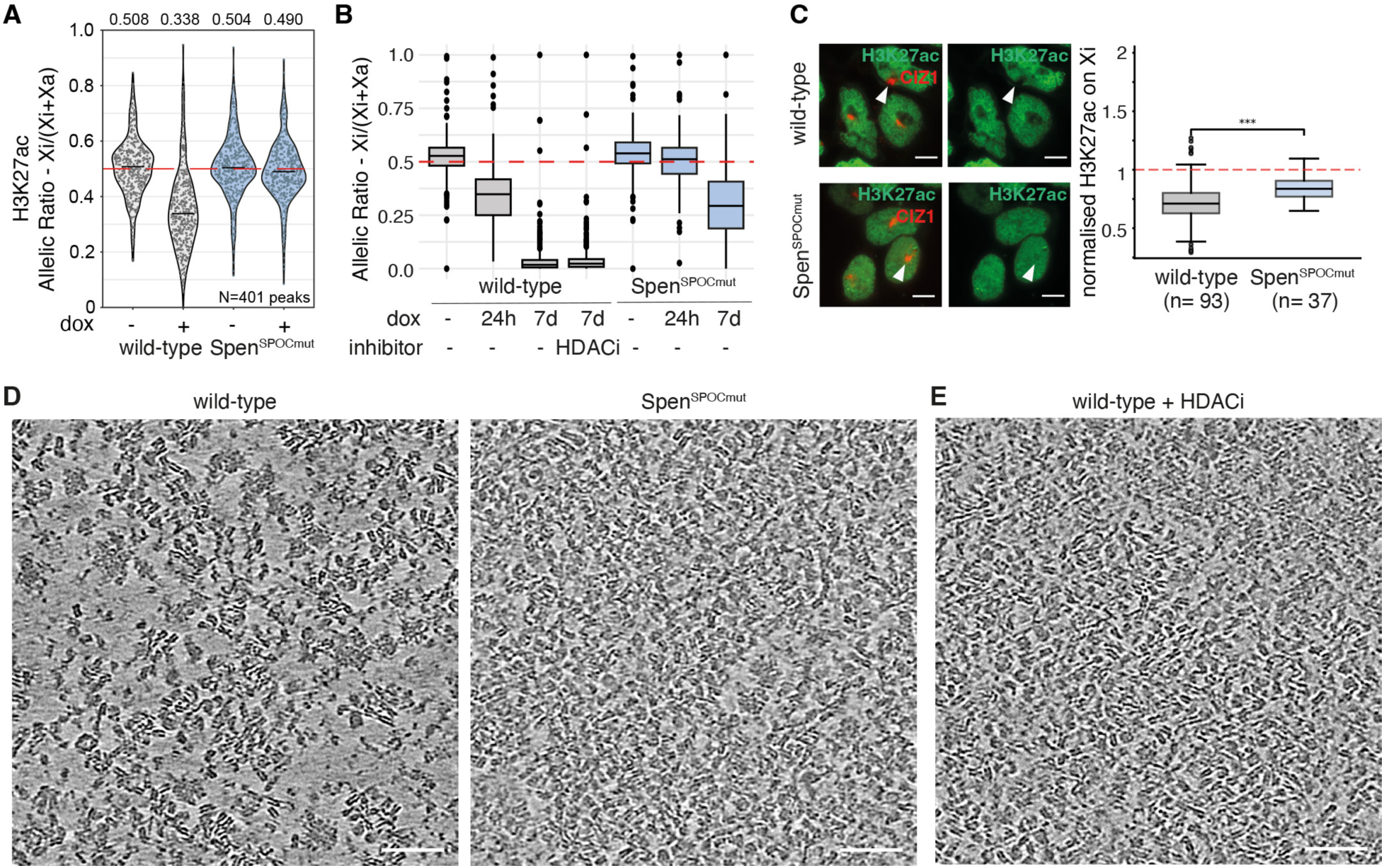
Histone deacetylation demarcates chromatin domains in the Xi. **(A)** A violin plot of allelic ratios of ChIP-seq reads overlapping H3K27ac peaks of Xa and Xi, showing the abrogation of Xist-mediated deacetylation of the Xi in SPEN^SPOCmut^. Points represent mean allelic ratios of individual peaks across two replicates and are horizontally jittered to fill violins. Horizontal bars indicate the median values of each sample, which are also shown numerically above. **(B)** A boxplot summarising allele-specific ChrRNA-seq analysis of X-linked gene expression in wild-type and Spen^SPOCmut^ cell lines. Cells without Xist induction (−dox) and with 24-hour Xist induction were maintained under ESC conditions, whereas cells with 7-day Xist induction were maintained under differentiation conditions. **(C)** Left: Immunofluorescence image of H3K27ac and CIZ1 (marking the Xi) in day-7 differentiated wild-type and Spen^SPOCmut^ mESCs; scale bar = 10 µm. Right: A boxplot showing H3K27ac signal on the Xi domain, normalised by the mean H3K27ac signal outside the Xi domain of each nucleus. P = 1.464e-05. **(D)** Comparison of cropped tomographic slices of wild-type and Spen^SPOCmut^ cells. For uncropped views, see Extended Data Fig. 5A and Supplementary Video 1 for wild-type; and Extended Data Fig. 8A and Supplementary Video 5 for Spen^SPOCmut^. **(E)** A cropped Xi tomographic slice from an HDACi-treated wild-type cell; scale bar = 50 nm. For uncropped views, see Extended Data Fig. 8B and Supplementary Video 6. Scale bars for **(D)** and **(E)** = 50 nm.

We went on to reconstruct cryo-tomograms of the acetylated Xi in day-7 differentiated Spen^SPOCmut^ mESCs using our established cryo-CLEM workflow. Interestingly, we observed that the acetylated Xi chromatin of Spen^SPOCmut^ cells had a distinct uniform distribution of nucleosomes and an associated reduction of chromatin-depleted regions (Fig. 3D, Extended Data Fig. 8A, and Supplementary Video 5). This contrasted with the hypoacetylated Xi, which showed noticeable chromatin-depleted regions demarcating chromatin domains (Fig. 3D, Fig. 1A, and Supplementary Video 1).

To ensure that the modified appearance of Xi chromatin in Spen^SPOCmut^ cells is linked to impaired histone deacetylation, we reconstructed cryo-tomograms of the Xi from day-7 wild-type cells treated with the HDAC class I inhibitor (HDACi) Romidepsin. As Romidepsin induces cell-cycle arrest and apoptosis^29^, we treated cells for only 24 hours, from day 6 of differentiation until vitrification on day-7. This 24-hour HDACi treatment was sufficient to increase the global level of histone acetylation (Extended Data Fig. 7E) but had no effect on X-linked gene silencing relative to day-7 wild-type cells (Fig. 3B). These conditions thus enabled us to discriminate the specific effect of histone acetylation on Xi chromatin structure independent of changes in Xi gene transcription. The native chromatin organisation of Xi in HDACi-treated wild-type cells (Fig. 3E and Supplementary Video 6) was remarkably similar to that observed in Spen^SPOCmut^ cells. This finding is consistent with optical microscopy studies showing that histone hyperacetylation decreases nuclear compartmentalisation^30^ and chromatin heterogeneity^31^, and results in more homogeneously distributed nucleosome signals^32^. A side-by-side comparison of full Xi tomograms from Spen^SPOCmut^ and HDACi-treated cells is shown in Extended Data Fig. 8A, B. Note that densities corresponding to putative RNPs located within chromatin-depleted regions are present in both instances, despite the relatively crowded distribution of the acetylated Xi nucleosomes.

## Multiscale homogenisation of densely packed chromatin on the acetylated Xi

To quantify chromatin heterogeneity in the native Xi under different conditions, we adapted a coefficient of variation (COV) approach, previously used to quantify chromatin heterogeneity by confocal microscopy^33^. We first measured local volume fraction occupied by nucleosomes within a unit volume of 28 × 28 × 28 nm^3^ across the whole tomogram, as depicted in a heat map (Fig. 4A). We then calculated the COV, defined as the ratio of the standard deviation (SD) of local volume fraction occupied by nucleosomes to its mean. The COV was lower in the acetylated Xi chromatin than in the hypoacetylated Xi chromatin (Fig. 4B), quantitatively confirming the reduced chromatin heterogeneity in the acetylated Xi chromatin.

**Figure 4.**
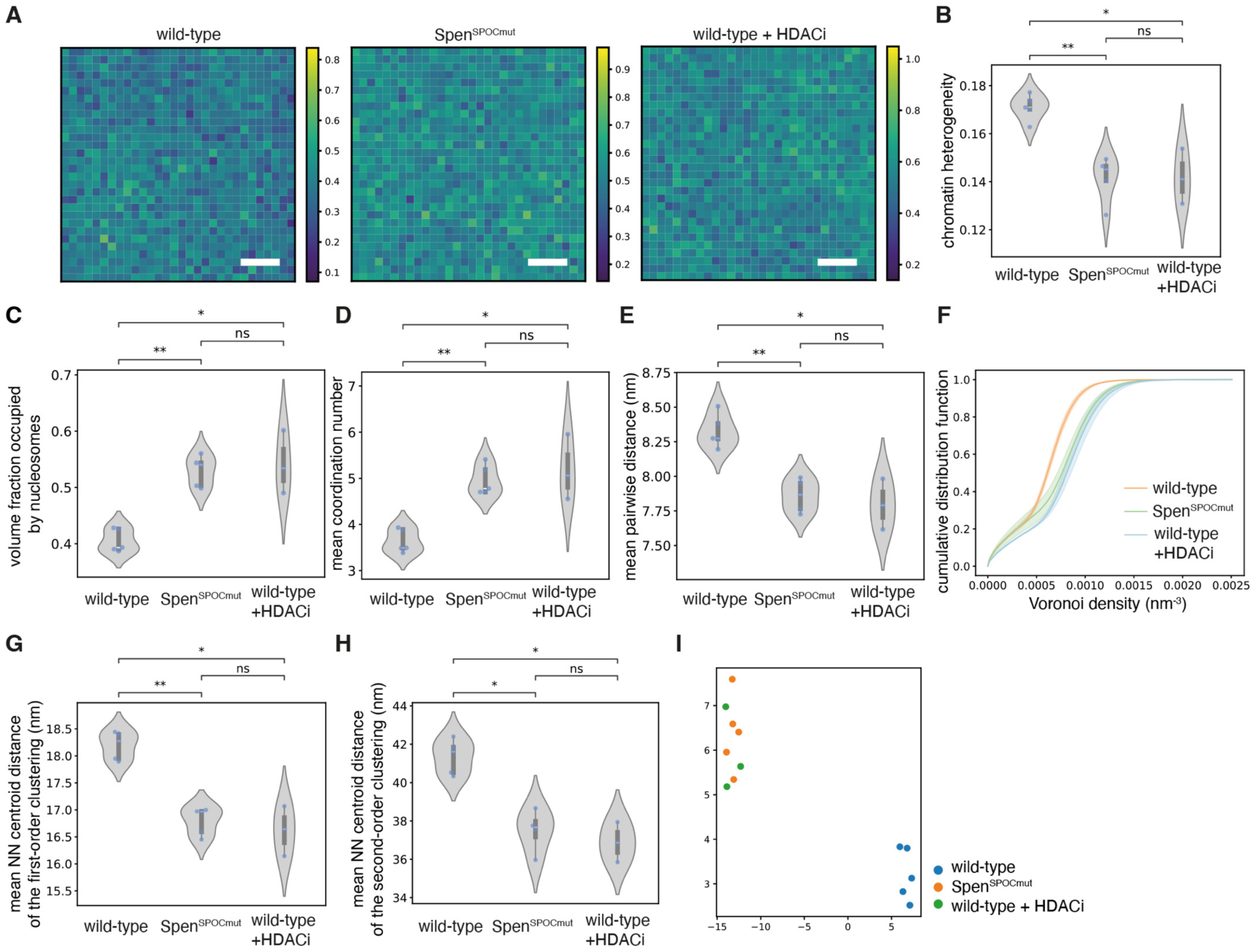
Histone acetylation in the Xi results in homogenisation and more densely packed chromatin organisation. **(A)** Heat maps representing local volume fraction occupied by nucleosomes in a unit volume (28 × 28 × 28 nm^3^) across the tomogram subvolume (839 × 839 × 28 nm^3^). The colour key for each heat map is provided on the right side of the heat map. Scale bar = 150 nm. Violin plots of **(B)** chromatin heterogeneity, calculated from the COV, **(C)** mean volume fraction occupied by nucleosomes, **(D)** mean coordination number of nucleosomes within a sphere of 12 nm radius, and **(E)** mean pairwise distances between NN nucleosomes. **(F)** A plot of the CDF of Voronoi-based local density. The line represents the mean, and the shaded region represents the standard error of the mean across tomograms. **(G and H)** Violin plots of mean pairwise distance between NN (**G**) first-order clusters and (**H**) second-order clusters. **(I)** UMAP of all quantitative chromatin-organisation features. Each dot represents a tomogram of the inactive X chromosome from wild-type, Spen^SPOCmut^, or HDACi-treated wild-type cells. **(B)-(E), (G)-(H)**: each dot represents a data point from a subvolume (839 × 839 × 46.6 nm^3^) of an independent tomogram, consisting of >20,000 extracted nucleosome particles. Exact P-values between all data points are provided in Supplementary Data Table 1. n = 5 (wild-type); 5 (Spen^SPOCmut^); 3 (wild-type + HDACi). (**F**) and (**I**) have the same n.

Next, capitalising on the direct visualisation of all nucleosomes in the Xi cryo-tomograms, we quantitatively examined the effect of histone acetylation on the hierarchical organisation of native Xi chromatin. First, consistent with the reduction in chromatin-depleted regions, the acetylated Xi chromatin of both Spen^SPOCmut^ and HDACi-treated wild-type cells showed higher mean volume fraction occupied by nucleosomes (Fig. 4C) and higher mean coordination number (Fig. 4D) than the hypoacetylated Xi chromatin of wild-type cells. In line with this, the acetylated Xi chromatin also showed shorter pairwise distances between NN nucleosomes (Fig. 4E) and higher Voronoi density (Fig. 4F) than the hypoacetylated Xi chromatin. At the nucleosome-clutch scale, obtained from first-order clustering, the acetylated Xi chromatin showed shorter spacing between NN nucleosome clutches (Fig. 4G, Extended Data Fig. 9A) and a higher number of clutches per unit tomogram volume (Extended Data Fig. 9B), compared with the hypoacetylated Xi chromatin. The same trend was observed at the chromatin domain level, obtained from second-order clustering (Fig. 4H, Extended Data Fig. 9C, D). STA map of the Xi nucleosome from Spen^SPOCmut^ cells showed extra density corresponding to histone H1, similar to wild-type Xi (Extended Data Fig. 9E). Collectively, these results quantitatively demonstrate that the acetylated Xi chromatin exhibits a more densely packed configuration than the hypoacetylated Xi chromatin, from the single nucleosome to the chromatin domain in unperturbed native cells.

Finally, we applied Uniform Manifold Approximation and Projection (UMAP)^34^ to these chromatin organisation features from individual tomograms of the Xi across distinct cell states. Tomograms from the same cell state clustered closely in the 2D embedding, and each cell state occupied a distinct region of the UMAP space (Fig. 4I). Importantly, tomograms representing the acetylated Xi from Spen^SPOCmut^ and HDACi-treated wild-type cells clustered together and remained well separated from the hypoacetylated wild-type Xi (Fig. 4I), indicating that histone acetylation is a major contributor to the observed chromatin features.

In summary, using genetic mutation and pharmacological inhibition, we directly visualised and quantitatively analysed the native Xi chromatin structure under distinct histone acetylation states. These results uncover the structural role of histone deacetylation in demarcating chromatin domains by promoting the formation of chromatin-depleted regions.

## Discussion

In this study, we have established and applied cryo-CLEM combined with 3D-targeted FIB milling to determine the native chromatin structure associated with a defined transcriptional state, facultative heterochromatin of the Xi in female mammalian cells. Optimised particle picking and STA analysis of nucleosomes facilitated downstream analysis of chromatin organisation across scales, from individual nucleosomes to chromatin domains. Direct visualisation of chromatin in unperturbed cells at sub-nucleosome resolution also enabled us to capture unknown heterogeneous densities in the chromatin-depleted regions in Xi tomograms which we interpret as putative RNPs. Given that gene transcription is largely shut down on the Xi of day-7 differentiated cells, it is likely that at least some of these densities, especially those surrounded by compact chromatin, represent Xist RNP molecules. However, at present we cannot rule out that putative Xi RNPs instead represent mRNPs derived from genes that escape XCI or from transcriptionally active regions of neighbouring chromosomes captured within individual Xi tomograms. In future work, approaches such as genetically encoded multimeric tagging^35^ or ferritin-based tagging^36^ will be required to unequivocally identify Xist RNPs in cryo-tomograms.

A striking observation from our analysis is that a substantial proportion of Xi nucleosomes bind histone H1, at least among those that contributed to reconstructing the well-defined STA map. Notably, the STA nucleosome map from other chromosomes lacks clear histone H1 density, implying that the H1-bound nucleosome population is not abundant, or that H1 binding is more dynamic, in other chromosomes^37^. Furthermore, mapping the 3D spatial distribution of histone H1-bound nucleosomes in the Xi of near-native cells reveals that they form the compact core of chromatin domains. In a recent study, H2AK119ub was proposed to enhance histone H1-mediated chromatin compaction in mESCs^38^. In addition, ablation of histone H1 was shown to lead to chromatin decompaction and derepression of Polycomb target genes in human cells, while H3K27me3 levels remained unchanged^39^. The presence of H1-bound nucleosomes in both day-5 and day-7 Xi chromatin is consistent with its direct link to Polycomb function, an early-stage Xi modification^5^. Our study thus presents structural evidence in support of a link between the Polycomb system and histone H1, and implicates this pathway in the compaction and silencing of facultative heterochromatin.

We observe that chromatin on the compacted Xi is organised into nanoscale domains, consistent with data obtained using super-resolution light microscopy^40^. We further observe multiscale compaction of Xi chromatin between day-5 and day-7 following the onset of Xist expression and mESC differentiation, as illustrated in the model in the left and centre panels of Figure 5. The timing of this switch implicates late-stage XCI pathways that act subsequent to the recruitment of the chromosomal protein SMCHD1 to the Xi^41^. These changes may be directly linked to SMCHD1 function in XCI, notably in relation to the long-range organisation and compaction of chromatin domains^42^. Other late-stage Xi chromatin modifications that could induce compaction include recruitment of the variant histone macroH2A and the establishment of CpG island DNA methylation at X-linked genes^5^. It is also possible that cumulative and/or combinatorial effects of early-stage modifications, specifically Polycomb/H1 at the core of nanoscale domains and the absence of acetylated nucleosomes, which are normally located at the surface of nanoscale domains^6,43^, contribute to progressive Xi chromatin compaction.

**Fig. 5.**
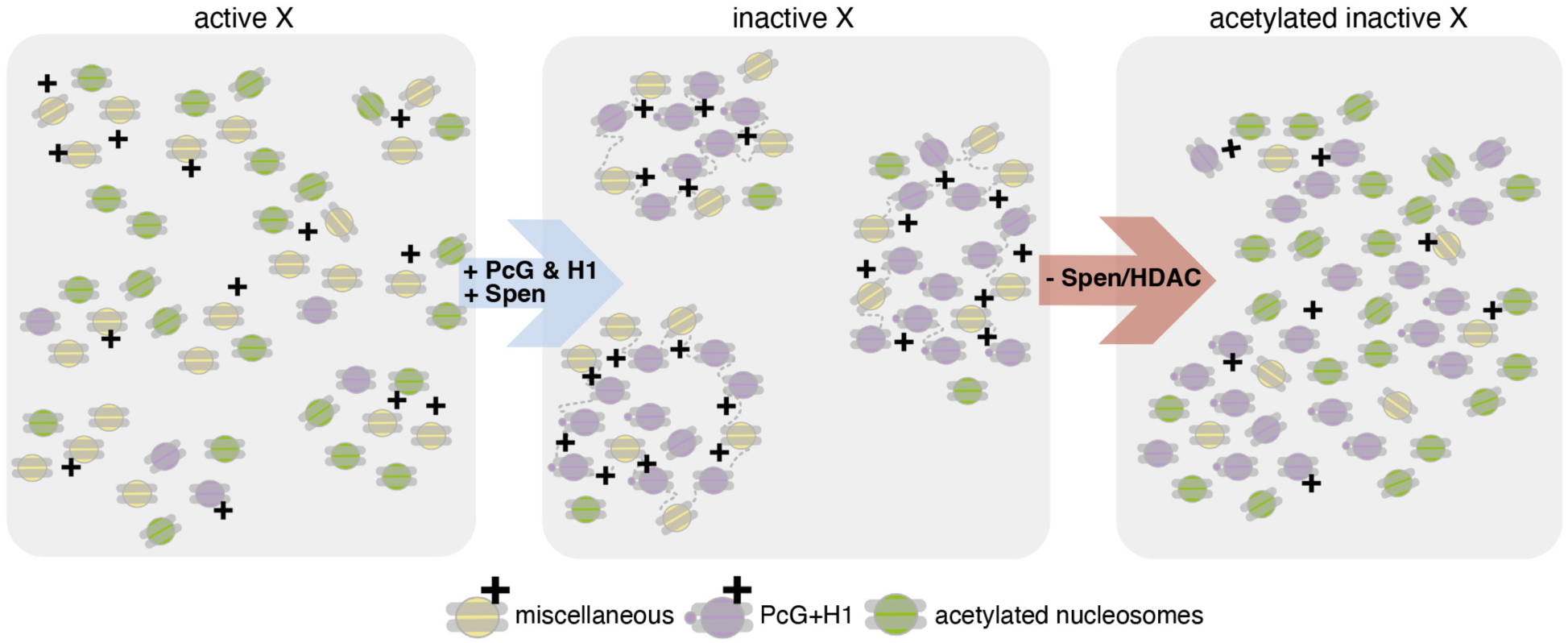
A model for Xi chromatin domain organisation and the role of histone acetylation. The left and centre schematics depict the transition of active X nanoscale chromatin domains to the configuration observed on the fully silenced Xi, indicating the inferred arrangement of positively charged miscellaneous and Polycomb group protein modified (PcG)/H1 nucleosomes, and charge-neutralised acetylated nucleosomes. The right panel depicts the changes that occur in the organisation of Xi chromatin domains in Spen^SPOCmut^ or HDACi-treated (−Spen/HDAC) cells.

The restoration of histone acetylation on Xi in Spen^SPOCmut^ or HDACi-treated cells results in homogenisation of nucleosomal distribution, with an associated reduction in the spacing between chromatin domains. This dramatic remodelling of chromatin domains results in individual nucleosomes being on average closer to one another, albeit in a uniform manner. A relatively dense packing of nucleosomes on the acetylated Xi seems counterintuitive in the context of evidence that histone tail acetylation inhibits the formation of compact 30-nm-like fibres in vitro^44^ and correlates with chromatin accessibility in cells^43^. Different factors likely underlie this finding. First, the acetylated Xi chromatin retains high levels of Polycomb/H1 and presumably other Xi-associated features, such as enrichment of SMCHD1. A second consideration is that histone acetylation is expected to alter the balance of interactions that drives chromatin to demix, rather than solely the intrinsic compaction of chromatin itself. Histone tails mediate both intramolecular contacts, with linker and nucleosomal DNA within the same fibre, and intermolecular contacts between fibres, and the partitioning between these two modes governs the connectivity, thermodynamic stability, and material properties of chromatin condensates^45,46^. Acetylation neutralises positively charged lysines on histone tails and weakens both classes of contact^47,48^. In this framework, the primary consequence of SPEN–HDAC3-mediated deacetylation is not solely local compaction but an increase in the contrast between chromatin-dense and chromatin-depleted regions, that is, the demarcation of domain boundaries reported here (Fig. 5, middle panel). Conversely, Xi acetylation lowers this contrast, the two-phase distribution collapses, and nucleosomes redistribute into volumes that were previously chromatin-depleted (Fig. 5, right panel). This interpretation is consistent with previous work demonstrating that histone acetylation dissolves chromatin droplets in vitro^49^ and reduces the growth of chromatin condensates, as well as chromatin heterogeneity in cells^33^.

In summary, our study advances understanding of the native chromatin structure of facultative heterochromatin, using the Xi as a model, at the chromatin-domain scale and below, a dimension previously concealed by the limitations of optical and sequence-mapping resolution. The cryo-CLEM strategy described herein provides a powerful new approach to further understand how different pathways contribute to the structure and organisation of chromatin on the Xi and at other sites in the genome.

## Methods

### Cell culture

Maintenance and monolayer differentiation of XX mESCs were performed as described previously^11^. To generate a wild-type XX mESC line tailored for cryo-CLEM, we derived a stable cell line expressing an mCherry-fused Ciz1 transgene from the Rosa26 locus in the established iXist-ChrX^1^^29^ mESC line^10^ using CRISPR/Cas9 homology directed repair (HDR). Briefly, iXist-ChrX^1^^29^ cells were co-transfected with an sgRNA/Cas9 plasmid targeting the Rosa26 locus and a mCherry-Ciz1 HDR plasmid using Lipofectamine 3000 (Thermo Fisher Scientific, TFS) according to the manufacturer’s protocol. Positively transfected cells were selected for using puromycin selection over 48 h (3.5 µg/ml) followed by ∼10 - 14 days growth until single mESCs colonies had formed. Single mESC clones were isolated and expanded for further testing.

To generate the Spen^SPOCmut^ mESC line, we performed CRISPR-assisted homologous recombination in the mCherry-Ciz1 iXist-ChrX mESC line described above, using the protocol and reagents from our previous study^15^. Both cell lines were validated using PCR, live imaging, and immunofluorescence (Extended Data Fig. 1B). We further confirmed the silencing function of the cell lines by performing chromatin RNA-seq as previously described^10^ (n = 1, Fig. 3B).

### ChIP-seq

Native ChIP-seq was performed largely as described in our previous study^10^, with all buffers supplemented with 5 mM of the deacetylase inhibitor sodium butyrate (Sigma). Detailed experimental procedure is provided in Supplementary Methods.

#### NGS read alignment and allelic split

ChIP-seq analysis was performed using previously established pipelines described in our previous study^15^. Fastq files were mapped using bowtie2 (v2.3.2) ^50^ to the 129S1xCast N-masked mm10 genome with parameters “–very-sensitive –no-discordant –no-mixed -X 2000”. Alignment files were then sorted by samtools and PCR duplicates were marked and discarded by picard-tools “MarkDuplicates” (Broad Institute). Aligned reads were assigned to separate files of CAST (‘genome1’) or Domesticus/129S1 (‘genome2’) genomes by SNPsplit (v0.2.0)^51^ using the “--paired” parameter and a strain-specific SNP file compiled from UCSC databases. Alignment (bam) files were sorted and indexed by samtools. Bigwig pileup tracks were generated by bamCoverage from deeptools, normalized by library size, and visualised with IGV (Broad Institute). Note that due to the previously described recombination in the iXist-ChrXDom line^10^, only the region of Chromosome X distal to the Xist locus is amenable to allelic analysis (‘ChrX1’).

#### Peak calling

Peak calling was performed on ‘unsplit’ (ie not allele-separated) bam files for each sample by MACS2 (v2.2.7.1)^52^ using parameters of “-f BAMPE -g mm --broad --broad-cutoff 0.05” compared to a background of sequencing input DNA. A consensus H3K27ac peak annotation was generated using a custom R script “K27acChIP_GeneratePeakList.R”, unifying regions covered by peaks in at least two sample replicates and filtering peaks by size between 50 bp and 10 kb. Peaks were also called from the ‘input’ DNA and subtracted from the consensus peak set as probable mapping artifacts.

#### Quantitative allelic analysis

The consensus H3K27ac peak set was parsed into gtf file format for featureCounts^53^, which was run with all allelic bam files to count reads overlapping peaks. These counts tables were used to calculate allelic ratios, defined as Xi/(Xi+Xa). Peaks which contained at least 10 allelically assigned reads in >80% of samples and showed biallelic signal in uninduced wild-type mESCs (0.15 < allelic ratio < 0.85) were used for analysis (n = 401 peaks on ChrX1).

### Cryo-EM grid sample preparation

Either wild-type or Spen^SPOCmut^ cells were cultured in a T25 flask under monolayer differentiation conditions for six days. On day 6 of differentiation, UltrAuFoil R2/2 200-mesh grids (Quantifoil) were glow-discharged for 1 min 20 s using the ’high’ setting on a Harrick plasma cleaner, and PBS supplemented with 0.1% gelatin was applied to the grids for 3–5 min in a 35 mm µ-dish (ibidi). The PBS with 0.1% gelatin was aspirated, and ∼ 200 µl of trypsinised differentiating cells at a density of 1 - 2 × 10⁵ cells/ml were subsequently plated directly onto the gelatin-coated grids. 5 to 10 min later, cells were inspected using an inverted microscope (10X objective lens) to confirm their settling and even distribution on the EM grid. Then, 1 ml of monolayer differentiation culture media was added to the dish. For wild-type cells to be treated with the HDAC inhibitor, 10 nM Romidepsin (Selleckem, S3020) was added at this point. Cells were cultured for one more day until vitrification on day-7. On day-7, the medium was changed to phenol-free Fluorobrite DMEM (TFS), and cell coverage on the grid and mCherry-CIZ1-bound Xi in the cells were validated using a live-cell imaging Olympus IX83 system fitted with a humidified chamber at 37 °C. Immediately prior to vitrification, grids were incubated with phenol-free Fluorobrite medium supplemented with 10% glycerol as a cryoprotectant for 3–10 min. Grids were blotted from the back side for 3–5 s at 37 °C and 80% humidity, plunge-frozen in liquid ethane using a Leica EM GP2 plunger, and stored in liquid nitrogen. Grids were clipped into standard autogrid rings, with an orientation mark added using a permanent marker to aid aligning the lamella orientation when the milled grids were transferred to a transmission electron microscope (TEM) (see below).

### 3D-targeted FIB milling of the Xi

The Xi was endogenously labelled by stably expressing mCherry-tagged CIZ1 in XX mESC lines with inducible Xist expression^10^. CIZ1 binds to Xist RNA, delineating the Xi territory at interphase^11^. We confirmed that mCherry-CIZ1 in our cell line was specific to the Xi by immunofluorescence (Extended Data Fig. 1B). The endogenous fluorescence signal was clear on cryo-light microscopy imaging (Extended Data Fig. 1A). In order to capture the Xi in a 100–150-nm-thick lamella from 5–10-µm-thick mESCs, we performed 3D-targeted FIB milling as follows:

1. Vitrified grids were sputter-coated with platinum (120 s), then coated with organometallic platinum (40 s) using the gas injection system (GIS) and finally sputter-coated with platinum again (120 s) in the Arctis (TFS). Next, using an argon (Ar) plasma focused ion beam (PFIB), "X" marks (4 µm in length) were generated near the cells, on the flat surface of the grid.
2. 3D stack data were collected in the reflective, red, and green fluorescence channels using the built-in fluorescence light microscopy module of the Arctis.
3. The coordinates of the Xi from mCherry-CIZ1 signal in the red fluorescence image (X1, Y1, Z1) were measured relative to the centre of the fiduciary "X" mark as (0, 0, 0). The coordinates of the target milling location in the FIB view (X2, Y2) were determined as follows: X2 = X1, Y2 = Y1 × cos 12° + Z1 × sin 12° (with a milling angle of 12°). The green channel was used to exclude any potential autofluorescence signal from the mESCs, which usually show fluorescence signal across all channels.
4. The top and bottom milling boxes centred on the target milling coordinates (X2, Y2) were drawn and semi-automatic milling was performed using autoTEM or webUI software (TFS), with a 30 kV Ar ion beam ranging from 2 nA to 60 pA.
5. When the lamella thickness reached approximately 200–300 nm, lamella were imaged using the red fluorescence channel to verify the presence of the Xi, and to determine the location for collecting tilt-series data by TEM.
6. Final polishing was performed manually using a 20 pA Ar ion beam, monitoring the intactness of the lamella in both SEM and FIB images.

### Cryo-electron tomography data acquisition

The autogrids with lamellae were loaded with the lamella orientation perpendicular to the tilt axis of the microscope, guided by the mark on the autogrid ring, or multi-specimen cassettes containing milled grids were transferred directly from the Arctis to the Titan Krios G4 (TFS), equipped with an Autoloader (TFS) and a Falcon 4i camera. Using Tomo5 software (TFS), we collected Atlas images across the whole grid, acquired low-magnification images of the targeted lamella, and overlaid these onto fluorescence images of the thinned lamella. Tilt-series data were acquired from the correlated fluorescence region for the Xi tomograms, or from a nearby nuclear region for other chromosomes, using a dose-symmetric scheme ^54^ spanning ± 58° from -12° (effectively 0° at the lamella plane), with 2–3° increments and a grouping of 3, at a calibrated pixel size of 2.33 Å at the specimen level, a defocus range of 2–4 µm, and a total dose of 100–150 e⁻/Å² in EER format, using a Selectris X energy filter with a 10 eV slit and a 70 µm objective aperture.

### Tilt-series data pre-processing and tomogram reconstruction

Tilt-series stacks were created after correcting for the gain reference, motion, and contrast transfer function (CTF) using WarpTools (--eer groupexposure=0.5)^55^. Tilt-series were aligned using AreTomo^56^ and tomograms were reconstructed using WarpTools^55^ at a binning factor of 4. Tomogram reconstruction was further polished using MissAlignment^57^. 3D denoising was performed on the tomograms for clear visualisation using Noise2Map in WarpTools (--dont_flatten_spectrum, --batchsize 12, -- dont_augment)^55^. Denoised tomograms were used to prepare Figures and Videos.

### Nucleosome particle picking

To pick nucleosome particles from the tomograms, we performed template matching using GAPSTOP software^58,59^ (Extended Data Fig. 2A). First, a single bin-4 raw tomogram of the Xi was used with a human nucleosome sub-tomogram averaging map as a template (EMD-16979). A constrained-cross correlation score map from the template-matching run was manually inspected by overlaying it onto tomograms in Napari^60^ to obtain an optimal threshold value for particle extraction, using cryoCAT^49,61^. Extracted particles were then visually inspected by overlaying them onto tomograms using the ArtiaX plugin^62^ in ChimeraX^63^. Nucleosome particles were extracted using WarpTools ts_export_particles (--2d)^55^ at bin 1, and 3D classification and refinement were performed using RELION-5^64^. Template matching on the entire dataset was carried out using the obtained STA nucleosome map as a template (Extended Data Fig. 2A). For this, template matching was performed on both the raw tomograms and the missing-wedge-predicted tomograms, obtained by applying the "refine" and "predict" steps using IsoNet2^65^ on the denoised tomograms. Visual inspection of the resulting cross-correlation score maps from the raw and missing-wedge-predicted tomograms showed general overlap, but each showed ∼20% unique particles picked at reasonable thresholding (Extended Data Fig. 2A).

A single threshold value across the whole tomogram resulted in uneven picking between more crowded and sparser regions. To improve the detection of particles in cryoCAT, we adapted a method of finding peaks employed in PyTME^66^, using a maximum filter with a kernel size comparable to the nucleosome particle^67^. Then a local mean and local SD were calculated in a defined cubic volume with input parameters. Different sets of parameters were tested manually to determine the optimal picking, which we visually inspected by overlaying the extracted particles at different parameters onto tomograms. Extracted particles from the raw and missing-wedge-predicted tomograms, obtained using the optimal parameters, were then combined into a final particle list using a script^68^ generated with the assistance of Claude (Anthropic, version: Sonnet 4.6), removing duplicates by excluding any particles that overlapped by > 10% of the nucleosome volume (Extended Data Fig. 2A).

### Nucleosome STA analysis

Final nucleosome particles were extracted using WarpTools (ver. 2.0.0) ts_export_particles (--2d)^55^ at bin 1. All 3D classification and 3D refinement jobs were performed as described below using RELION-5^64^. To further improve the alignment of particles picked by template matching, we first performed 3D classification using local angular searches only (number of classes: 1; regularisation parameter T: 4; mask diameter: 270 Å) using the STA nucleosome map (--lowpass 50 Å) obtained from one of the tomograms (described above) as a reference map. The output particle star file from this job was then used to perform all downstream 3D classification (number of classes: 6; regularisation parameter T: 4; mask diameter: 270 Å) using differently modified nucleosome reference maps (--lowpass 30 Å). Reference maps used were: chromatosome, EMD-37149; and unmodified nucleosome, EMD-70743. We selected the best class from each and performed 3D refinement using solvent-flattened FSCs and blush regularisation. Finally, further refinement was performed using M refine^69^. The filtered map from the M refine job was used to prepare STA map Figures.

### Contextual analysis of nucleosomes and chromatin domains

Using the nucleosome coordinates and orientation from the final particle list, we calculated a list of feature-descriptors as follows.

#### Feature-descriptors at individual nucleosome level

For each nucleosome, we calculated the following feature-descriptors (N1 – N6):

N1. Pairwise distance between a nucleosome and its NN nucleosome.
N2. Pairwise distance between the nucleosome and its 8th, 27th, 64th, 125th, 216th, 343rd, 512th, 729th, 1000th NN nucleosome.
N3. Using the approach described in a previous report^46^, the angle 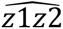 between a nucleosome’s Z-axis vector *z*_1_ and the Z-axis vector of its NN nucleosome, *z*_2_. Here, the Z-axis of the nucleosome was defined as the direction along which the nucleosome projection is maximum (i.e. perpendicular to the nucleosome plane).
N4. The angle 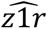 between a nucleosome’s Z axis vector and the vector *r*, in which starts from the centre of the nucleosome and ends at the centre of the NN nucleosome.
N5. Using the approach described in a previous report^46^, we categorize each nucleosome to 3 orientations, face-to-face, side-to-side and face-to-side with a slight variation as follows:
If 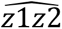 ∈ [30°, 150°] then we categorize the nucleosome as “face-to-side”
If 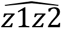 ∉ [30°, 150°] then if 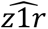 ∈ [30°, 150°] then we categorize the nucleosome as “side-to-side” and if 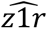 ∉ [30°, 150°] we categorize the nucleosome as “face-to-face”.
N6. Coordination number. We measured the number of neighbouring nucleosomes within different spheres of specific radii: 12 nm, 24 nm, 36 nm, 48 nm, 60 nm, 72 nm, 84 nm, 96 nm, 108 nm, 120 nm, and 132 nm.

#### Global feature-descriptors for the entire tomogram area used for nucleosome particle extraction

For each tomogram, we calculated the following features-descriptors for the set of nucleosomes (SN1 – SN6):

SN1. The fraction of volume occupied by nucleosomes in the volume of the entire tomogram used to detect nucleosomes. The total volume occupied by nucleosomes was calculated approximately as the number of nucleosomes times the average volume of a single nucleosome (550 nm^3^), divided by the total volume of tomogram we used for template matching.
SN2. The fraction of volume occupied by nucleosomes in the subvolume of the tomogram used to detect nucleosomes (120 × 120 × 120 pixel area of the tomogram at binning 1). We generated a heatmap representing the fraction of volume occupied by nucleosome in this subvolume. We calculated the mean and SD.
SN3. We calculated the COV, the ratio of SD and mean from SN2.
SN4. We calculated the mean and SD of for N1, N2 and N6.
SN5. We calculated constrained versions of the SN4 values where only nucleosomes that have their NN within 12 nm were considered to calculate the mean and SD.
SN6.

a. We generated histograms for the N1. For each we used the FITTER^70^ software to find the top 5 best matching distributions after each had its parameters set to best fit the histogram. The metric used to compare each distribution’s fitting is the sum square error.
b. After performing this on many histograms we detected that for N1, the 2 most commonly matching distributions were the Rayleigh and Gamma distribution. The parameters selected to best fit the Rayleigh and the Gamma distribution to the histogram of N1 were used as features-descriptors.
c. We calculated the Pair Distribution Function. We also calculated the Normalized Pair Distribution Function, using a random distribution of an equal number of nucleosomes in a volume of the same size as the area of the tomogram used to detect nucleosomes.

#### Feature-descriptors at first-order and second-order cluster level

We clustered nucleosomes using the Hierarchical Density-Based Spatial Clustering of Applications with Noise (H-DBSCAN)^20^. H-DBSCAN has 2 primary parameters to select. The Minimum Cluster Size (MCS) is the minimum number of nucleosomes in a group for it to be considered a cluster. The minimum number of samples (MNS) in a neighbourhood (including itself) is the minimum number for a point to be considered as a core point and added to a cluster. We used 4 for the MCS and 2 for the MNS.

For each cluster we calculated the following features-descriptors (C1 – C5):

C1. Population of nucleosomes within a cluster
C2. The coordinates of centroid of each cluster
C3. The mean pairwise distance of NN nucleosomes within a cluster
C4. The SD of the pair-wise distance of NN nucleosomes within a cluster
C5. Mean pairwise distance of NN clusters based on their centroid coordinates

For each tomogram, we calculated the following features-descriptors for the set of clusters (SC1 – SC3):

SC1. The mean and SD of the population of this set of clusters. (The number of nucleosomes counted as noise and were not assigned to a cluster)
SC2. The mean and SD of pairwise distance between a nucleosome and its NN (N1) for all nucleosomes that have not been categorized as noise.
SC3. The mean and SD of the pairwise distance of NN clusters based on their centroid coordinates

#### Utilization of feature-descriptors and dimensionality reduction

For each combination of sample (e.g., day-5 and day-7 differentiated wild-type) and gene activity state (e.g., Xi or other chromosomes), we used multiple tomograms to generate feature-descriptors (each N was annotated for each Figure). We obtained the P-value of this comparison using the Mann-Whitney U test^71^. Throughout the paper, P-values are annotated as follows: ns, 5.00 × 10⁻² < P ≤ 1.00; *, 1.00 × 10⁻² < P ≤ 5.00 × 10⁻²; **, 1.00 × 10⁻³ < P ≤ 1.00 × 10⁻². Exact P-values for each comparison are reported in Supplementary Data Table 1.

We utilized 36 nucleosome and clustering level feature-descriptors to describe each tomogram as a point in a 36-dimensional space. Using UMAP, we reduced the number of dimensions to 2. We used 5 neighbours and minimum distance of 0.1 to produce the figure 4I. We used Euclidean distances as a distance metric.

## Data Availability

We have deposited Cryo-ET maps at the Electron Microsocpy Data bank (EMDB) under accession numbers EMD-59329 (chromatosome of the Xi, day-7 wild-type), EMD-59330 (chromatosome of the Xi, day-5 wild-type), EMD-59331 (unmodified nucleosome of other chromosomes, day-7 wild-type), EMD-59292 (chromatosome of the Xi, Spen^SPOCmut^). Raw tilt-series data and reconstructed tomogram were deposited in EMPIAR under accession number EMPIAR-13877. ChIP-seq data were deposited in GEO accession GSE344252.

## Code availability

Our code is available at Zenodo^67,68,72^.

## Acknowledgments

We would like to thank Peijun Zhang and James Gilchrist (eBIC, Diamond Light Source) for help and support with establishing the cryo-CLEM workflow, Rosana Collepardo Guevara, Maria Julia Maristany and Jan Huertas Martin (Dept. of Chemistry, University of Cambridge, UK) for critical reading and input on biophysical aspects of chromatin organisation, and Laura Shemilt (Rosalind Franklin Institute, Harwell, UK) for help and guidance on the production of software and infrastructure optimization. This work was funded by grants to NB from the Wellcome Trust (215513/Z/19/Z), UKRI (EP/Y029062/1) and Leverhulme Trust (RPG-2025-241), and a Marie Skłodowska-Curie Individual Fellowship (797230) and Instruct-eric grant (PID-38380,37102) awarded to JC. Microscopy experiments were conducted at the OPIC electron microscopy facility, an Instruct centre, founded and supported by Wellcome and MRC.

## Author contributions

J.C. and N.B. conceptualised the project. J.C. and A.C. generated cell lines. J.C. and J.B. performed genomic analyses. J.C. established and performed the cryo-CLEM workflow and data processing. J.C. and D.B. developed code. J.C. and N.B. wrote manuscript with input from D.B., J.B., and A.C..

## Competing interests

Authors declare that they have no competing interests.

---

**Extended Data Figure 1.**
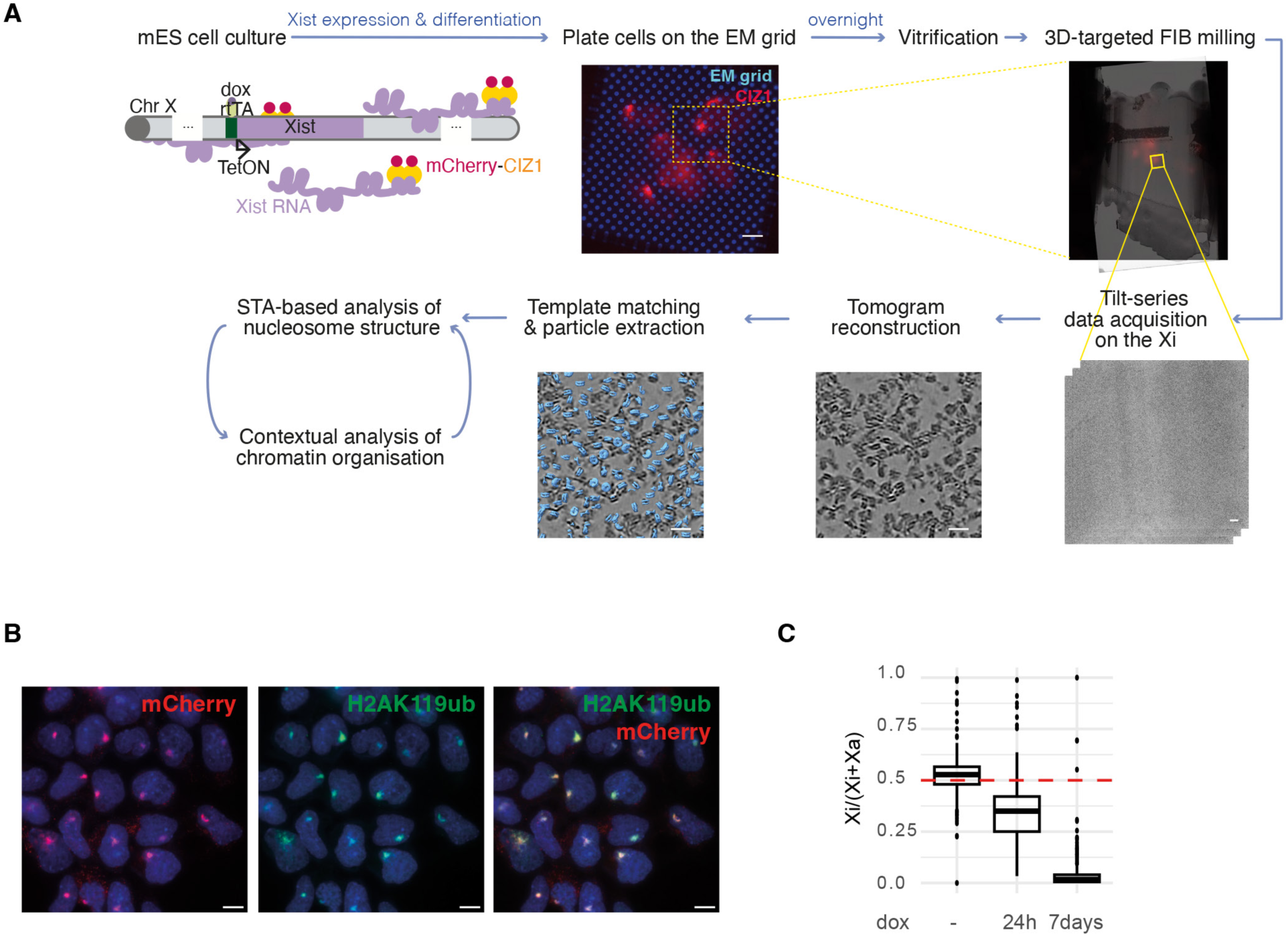
**(A)** A schematic and representative images describing the comprehensive workflow for analysing the in situ chromatin structure of the Xi. Scale bar = 10 µm for the light microscopy image; 50 nm for the tilt-series data image; 20 nm for the cropped tomogram slices. rtTA: reverse tetracycline-controlled transactivator, TetON: Tet-On promoter. **(B)** IF of day-7 differentiated mCherry-CIZ1-expressing female mESCs. The nucleus was stained using DAPI. Scale bar = 10 µm. **(C)** A boxplot summarising allele-specific ChrRNA-seq analysis of X-linked gene expression in wild-type cells. Cells without Xist induction (−dox) and with 24-hour Xist induction were maintained under ESC conditions, whereas cells with 7-day Xist induction were maintained under differentiation conditions.

**Extended Data Figure 2.**
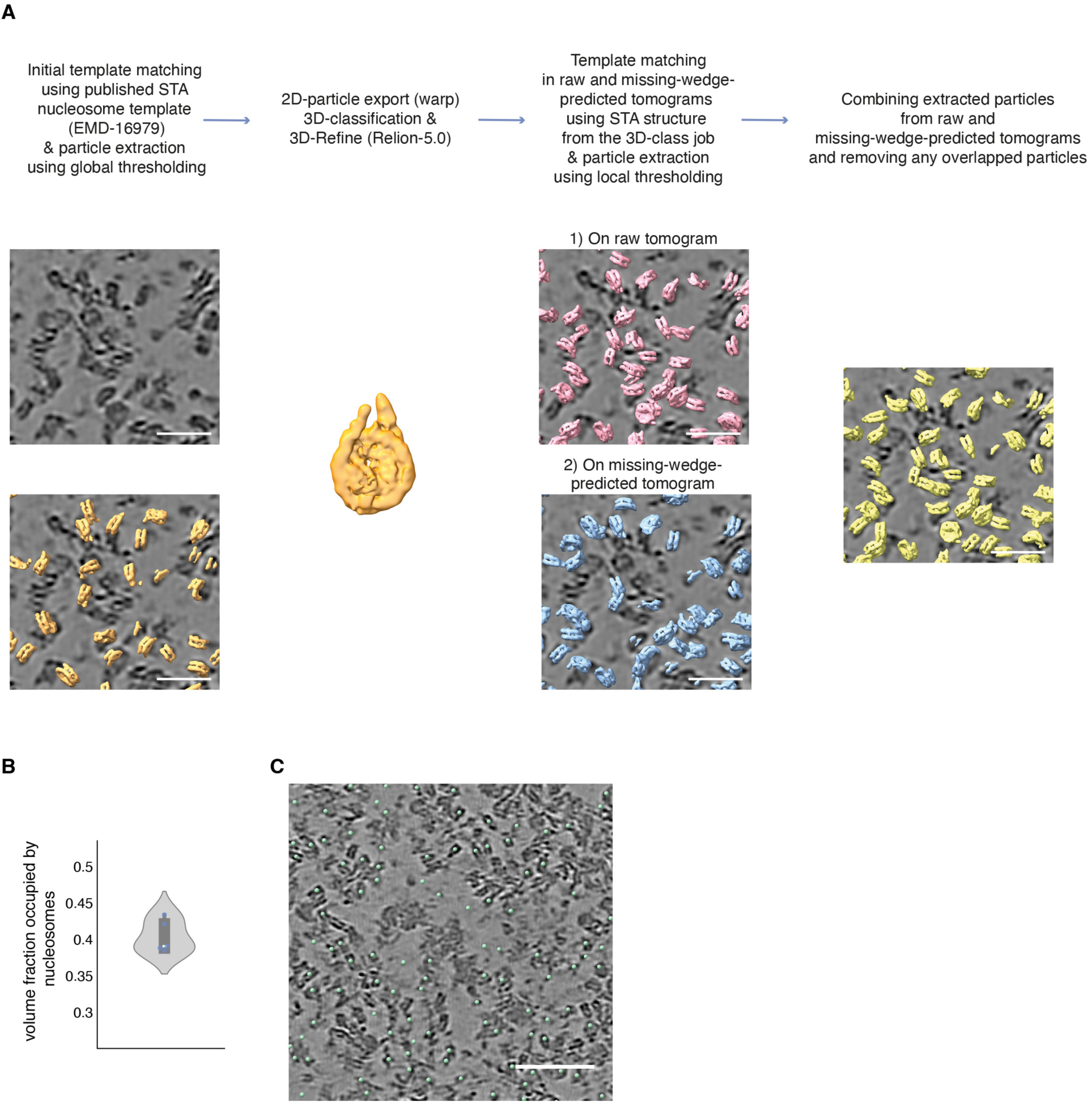
**(A)** A workflow for optimised template matching and particle extraction. Scale bar = 25 nm. **(B)** A plot of volume fraction occupied by nucleosomes within the tomogram volume across 5 Xi tomograms. **(C)** A cropped tomogram slice (equivalent to Fig. 1C) with nucleosomes picked by template matching overlaid as green spheres. Scale bar = 50 nm.

**Extended Data Figure 3.**
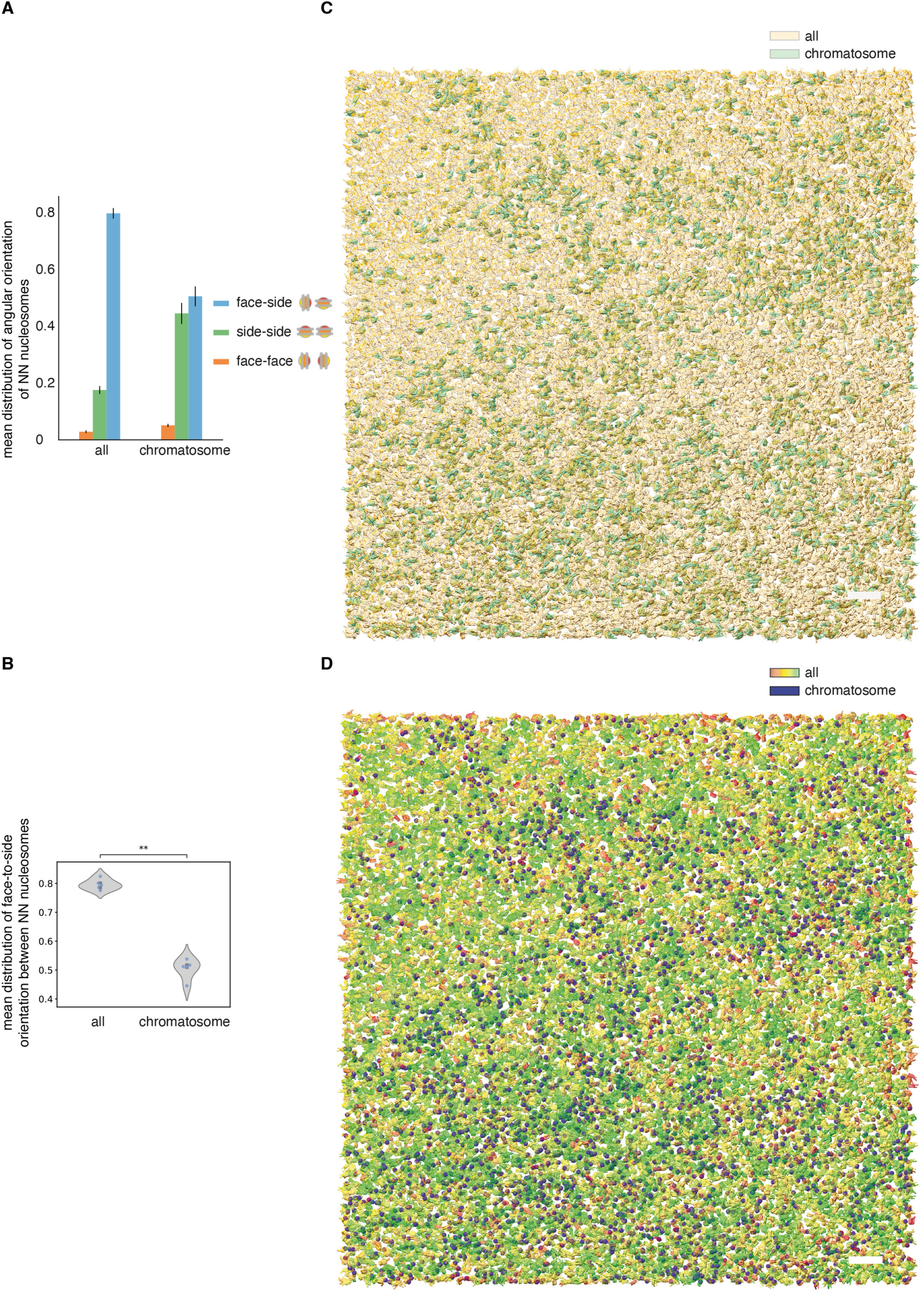
**(A)** A bar plot of the mean distribution of pairwise angular orientation between NN nucleosome pairs within a constraint sphere of radius 12 nm, for all nucleosomes or chromatosomes, in tomograms (n = 5). Error bars represent SD. For the detailed definition of each orientation group, see Methods. **(B)** A violin plot showing the mean distribution of face-to-side orientation only, from (**A**) (n = 5, P = 0.0079). **(C)** Extracted chromatosome (green) and all-nucleosome (orange) particles are mapped back to the 3D tomogram volume (z = 35 nm). Scale bar = 50 nm. **(D)** All nucleosomes are depicted as surface representations, and chromatosomes as purple spheres. Both are mapped within the 3D tomogram volume (z = 35 nm). All nucleosomes are coloured according to the number of neighbouring nucleosomes per nucleosome within a sphere of 12 nm radius (i.e., coordination number), from red (low) to green (high). Scale bar = 50 nm.

**Extended Data Figure 4.**
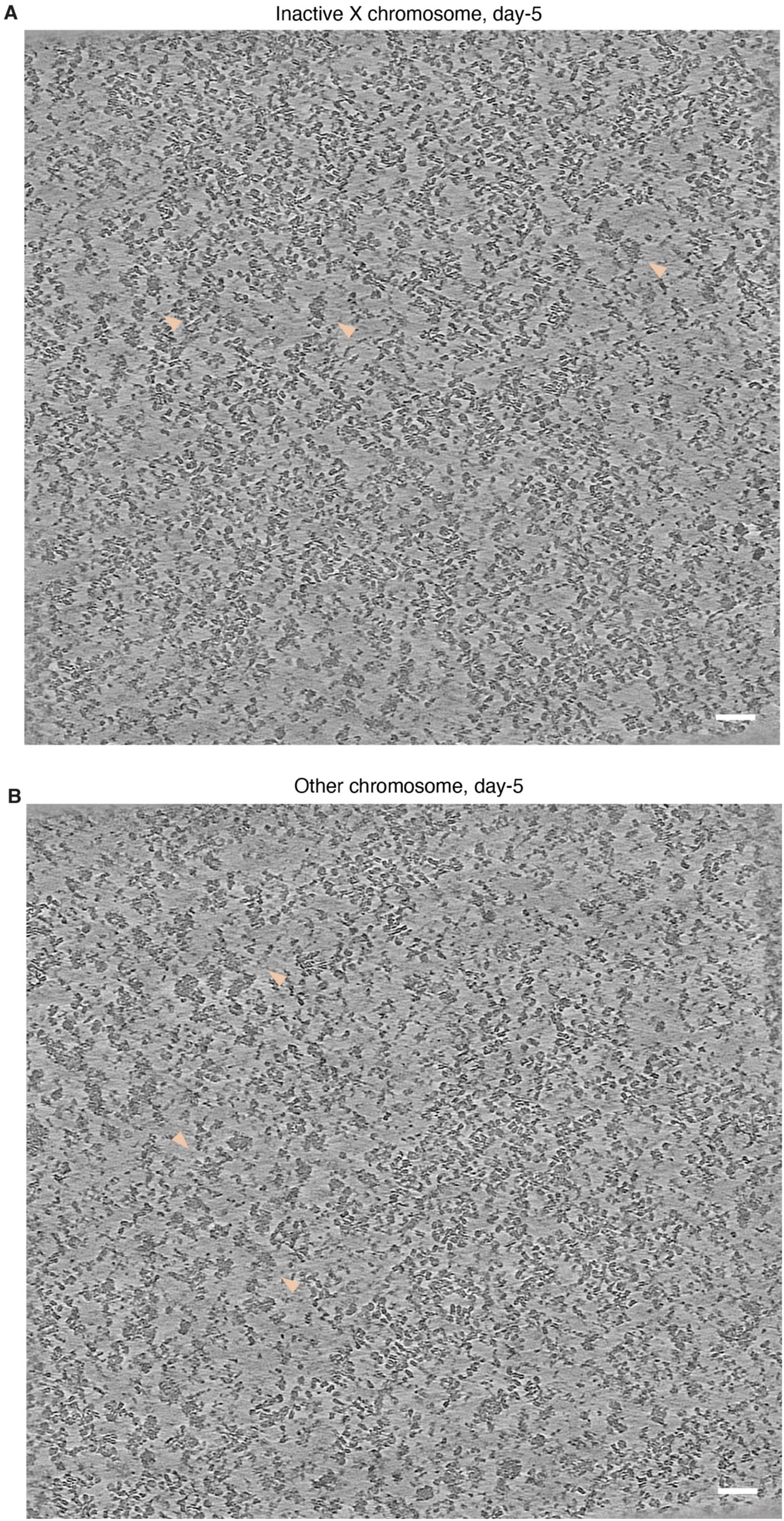
Tomogram slices of **(A)** the Xi and **(B)** other chromosomes from day-5 differentiated cells. Pink arrows indicate putative RNPs residing in chromatin-depleted regions. Scale bar = 50 nm.

**Extended Data Figure 5.**
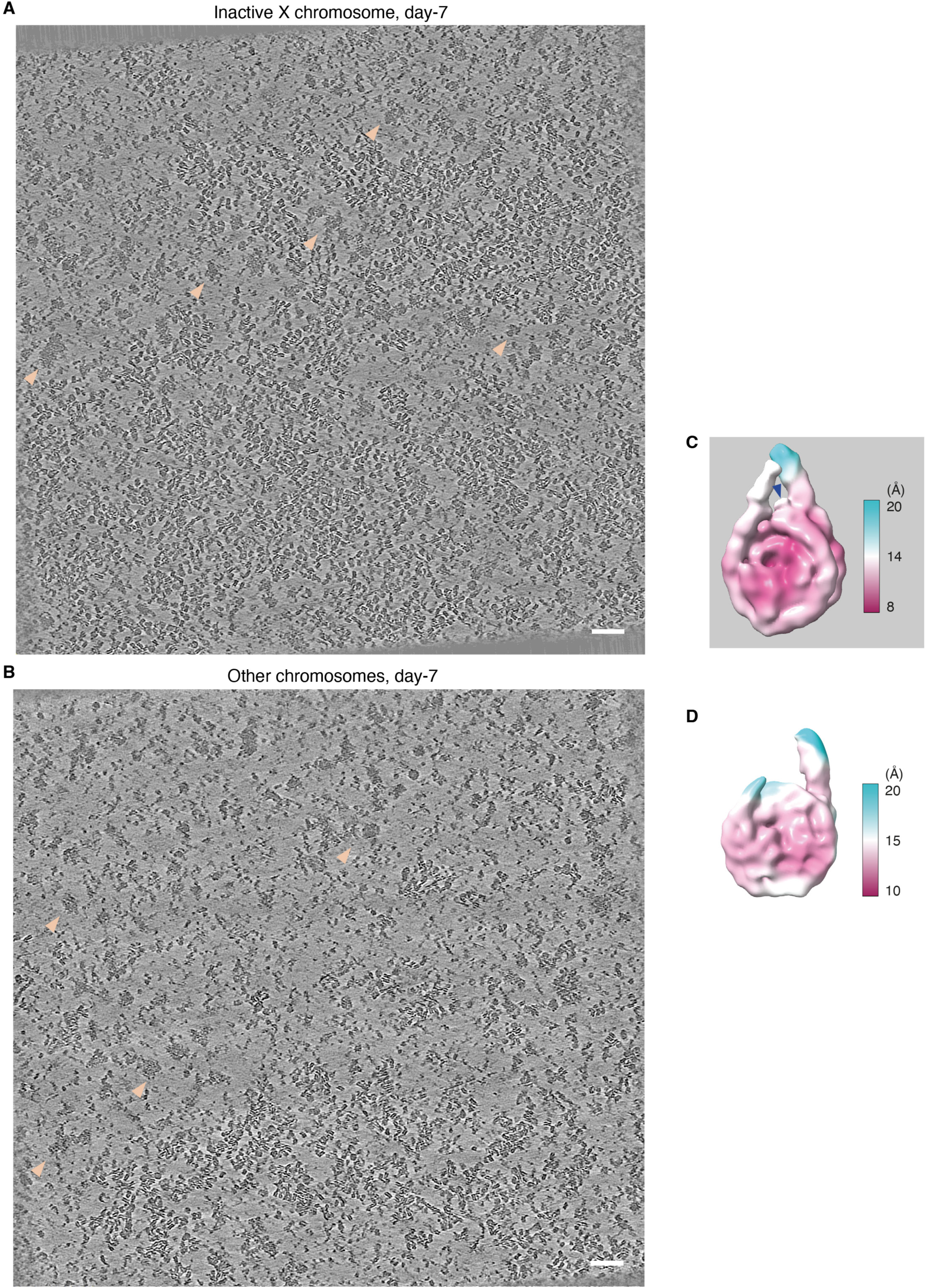
Tomogram slices of **(A)** Xi and **(B)** other chromosomes from day-7 differentiated cells. Pink arrows indicate putative RNPs residing in chromatin-depleted regions. Scale bar = 50 nm. **(C)** STA map of the Xi chromatosome from day-5 differentiated cells, coloured by local resolution. Arrow indicates putative density of histone H1. **(D)** STA map of the nucleosome from other chromosomes from day-7 differentiated cells, coloured by local resolution.

**Extended Data Figure 6.**
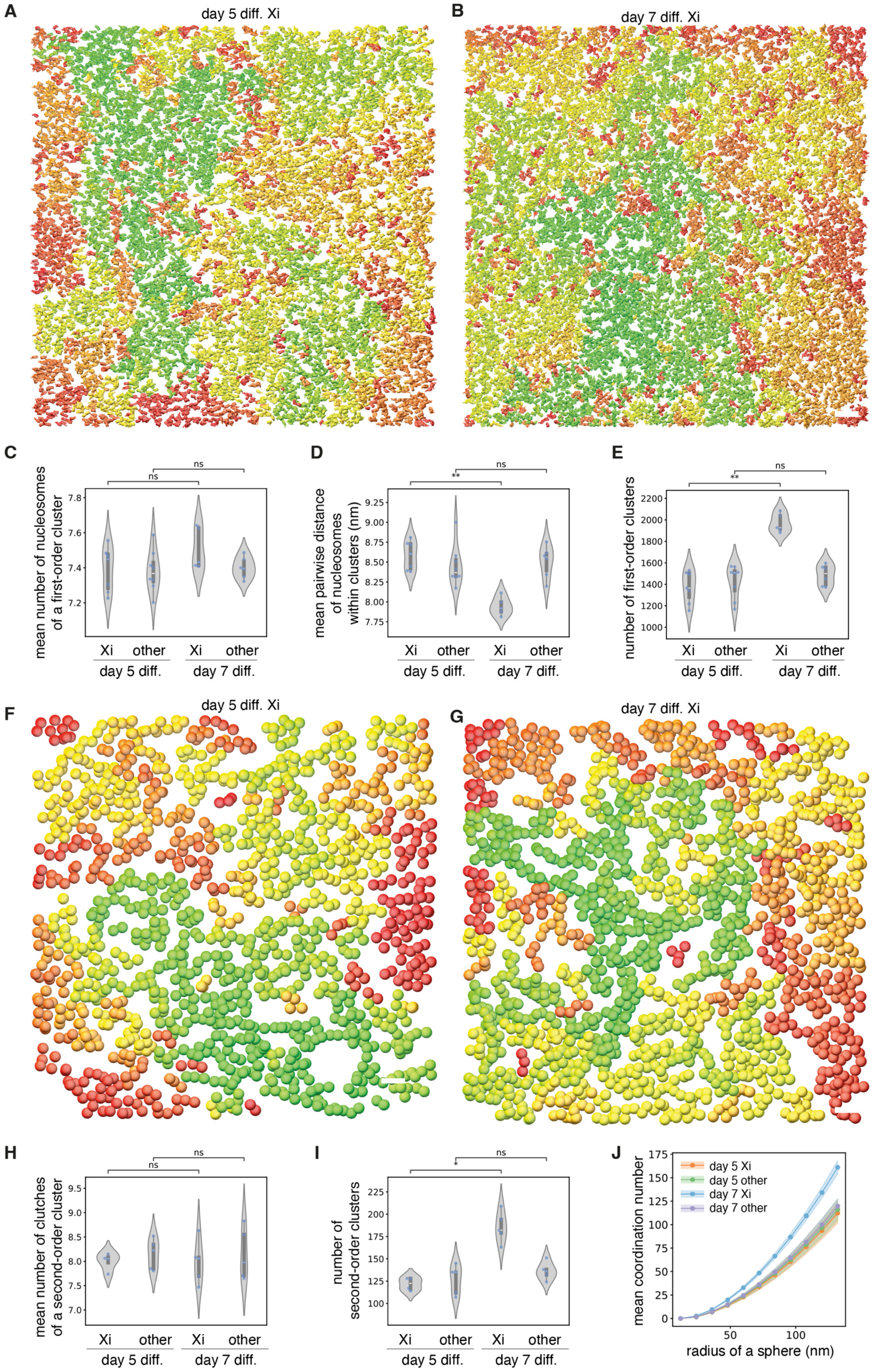
**(A, B)** Nucleosome clustering in day-5 (**A**, corresponding to Extended Data Fig. 4A, Supplementary Video 2) and day-7 differentiated Xi (**B**, corresponding to Extended Data Fig. 5A, Supplementary Video 1) within the 3D tomogram volume (z = 46.6 nm) using H-DBSCAN. Nucleosomes belonging to the same cluster are depicted in the same colour shade. Scale bar = 50 nm. Violin plots of **(C)** the mean number of nucleosomes in a first-order cluster, **(D)** the mean pairwise distance between NN nucleosomes within first-order clusters, and **(E)** the number of first-order clusters within the tomogram volume. **(F, G)** Second-order clustering of first-order clusters of the day-5 Xi (**F**, corresponding to Extended Data Fig. 4A, Supplementary Video 2) and day-7 Xi (**G**, corresponding to Extended Data Fig. 5A, Supplementary Video 1) within the 3D tomogram volume (z = 46.6 nm) using H-DBSCAN. Each first-order cluster is depicted as a sphere with a diameter of 28 nm around the centroid of the cluster, for simplification. First-order clusters belonging to the same second-order cluster are depicted in the same colour shade. Scale bar = 50 nm. Violin plots of **(H)** the mean number of nucleosome clutches in a second-order cluster and **(I)** the number of second-order clusters within the tomogram volume. **(J)** A plot of the mean coordination number of nucleosomes within spheres of increasing radius. Solid lines represent the mean value obtained from the tomograms of each category, and shading represents the SD. **(C)-(E), (H)-(I)**: each dot represents a data point from a subvolume (839 × 839 × 46.6 nm^3^) of an independent tomogram (n = 7 (day-5, Xi); 8 (day-5, other); 5 (day-7, Xi); 6 (day-7, other)), consisting of >20,000 extracted nucleosome particles. P values are annotated as follows: ns, 5.00 × 10⁻² < P ≤ 1.00; *, 1.00 × 10⁻² < P ≤ 5.00 × 10⁻²; **, 1.00 × 10⁻³ < P ≤ 1.00 × 10⁻². Exact P values for each comparison are reported in Supplementary Data Table 1. (**J**) uses the same n.

**Extended Data Figure 7.**
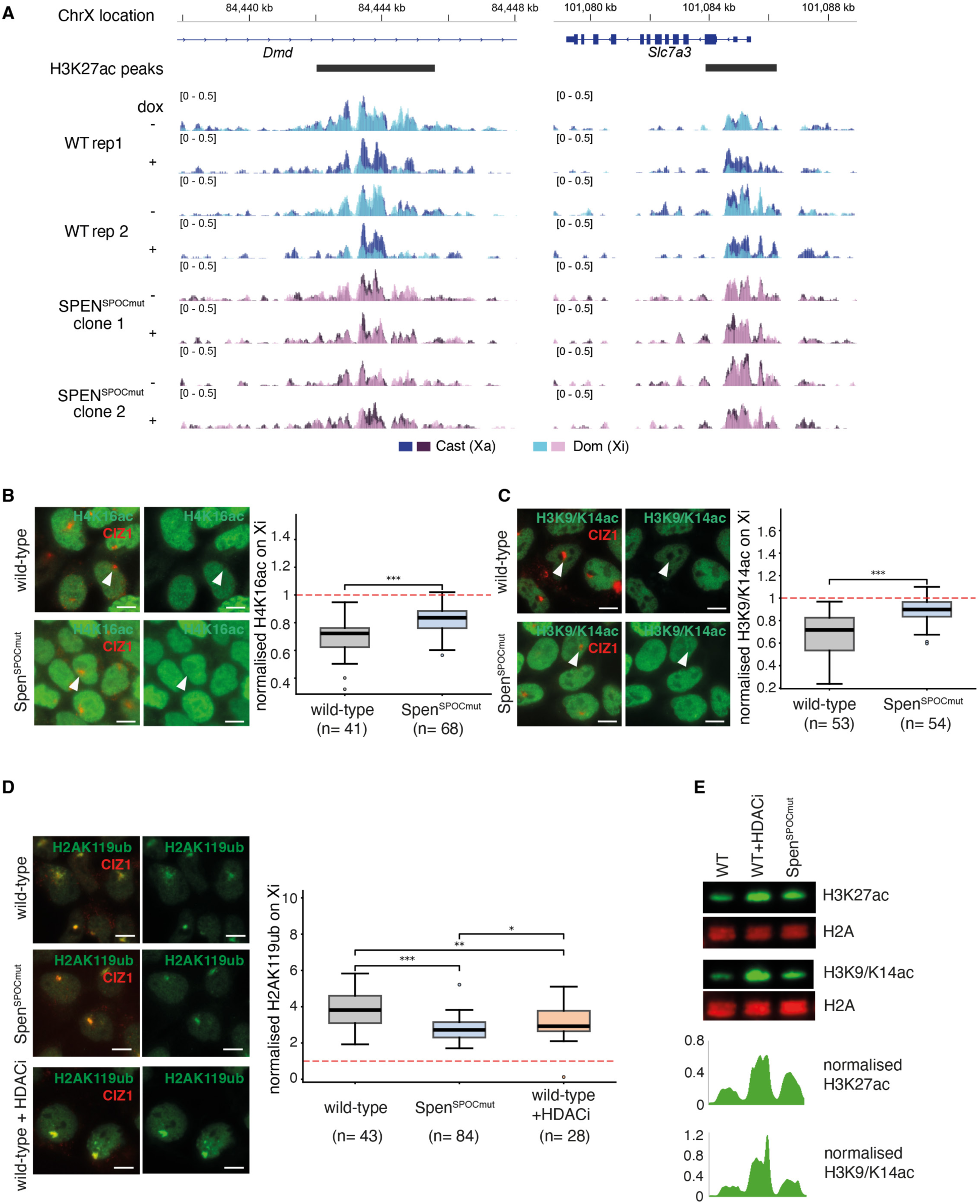
**(A)** Genome-browser (IGV) tracks of allele-specific H3K27ac ChIP-seq at two example peak regions: an intronic enhancer in the *Dmd* gene and the *Slc7a3* gene promoter. Deacetylation of the Xi allele is evident after 24 hours of Xist induction in wild-type cells, and is abolished in the Spen^SPOCmut^ cells. **(B)** Left: IF image of H4K16ac and CIZ1 (marking the Xi) in day-7 differentiated wild-type and Spen^SPOCmut^ mESCs; scale bar = 10 µm. Right: A boxplot showing H4K16ac signal on the Xi domain, normalised by the mean H4K16ac signal outside the Xi domain of each nucleus. P = 6.976e-06. **(C)** Left: IF image of H3K9/K14ac and CIZ1 in day-7 differentiated wild-type and Spen^SPOCmut^ mESCs; scale bar = 10 µm. Right: A boxplot showing H3K9/K14ac signal on the Xi domain, normalised by the mean H3K9/K14ac signal outside the Xi domain of each nucleus. P = 3.44e-10. **(D)** Left: IF image of H2AK119ub and CIZ1 in day-7 differentiated wild-type, Spen^SPOCmut^, and HDACi-treated wild-type mESCs; scale bar = 10 µm. Right: A boxplot showing H2AK119ub signal on the Xi domain, normalised by the mean H2AK119ub signal outside the Xi domain of each nucleus. P = 1.247e-08 (WT vs Spen^SPOCmut^); 0.008257 (WT vs HDACi); 0.02026 (Spen^SPOCmut^ vs HDACi). **(E)** Western blot of H3K27ac and H3K9/K14ac in the indicated cells. H2A was used as a loading control. The graphs below show the quantification of signal intensity normalised against the loading control (H2A).

**Extended Data Figure 8.**
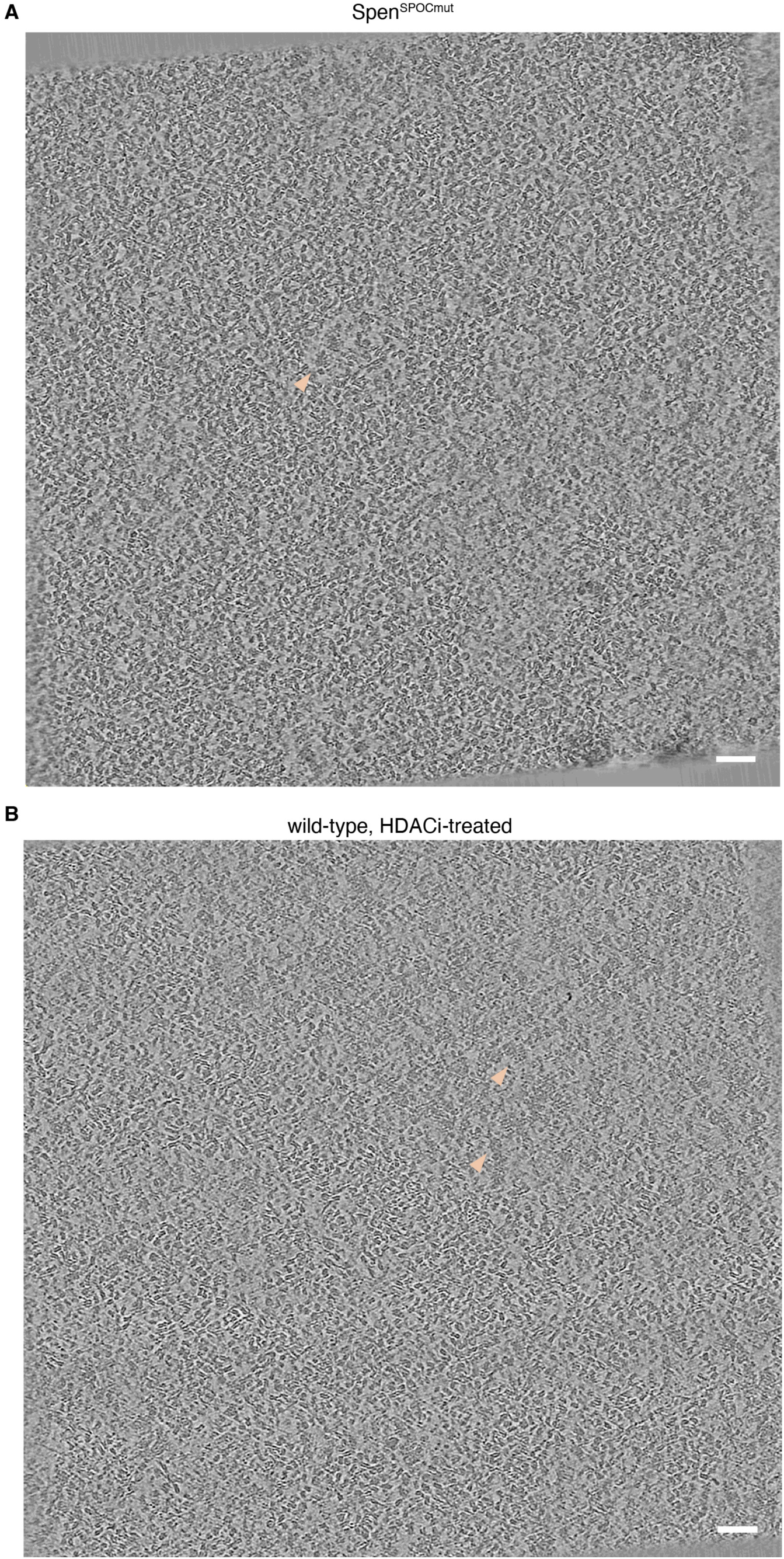
Xi tomogram slices from **(A)** Spen^SPOCmut^ and **(B)** HDACi-treated wild-type cells. Scale bar = 50 nm. Pink arrows indicate potential RNP densities.

**Extended Data Figure 9.**
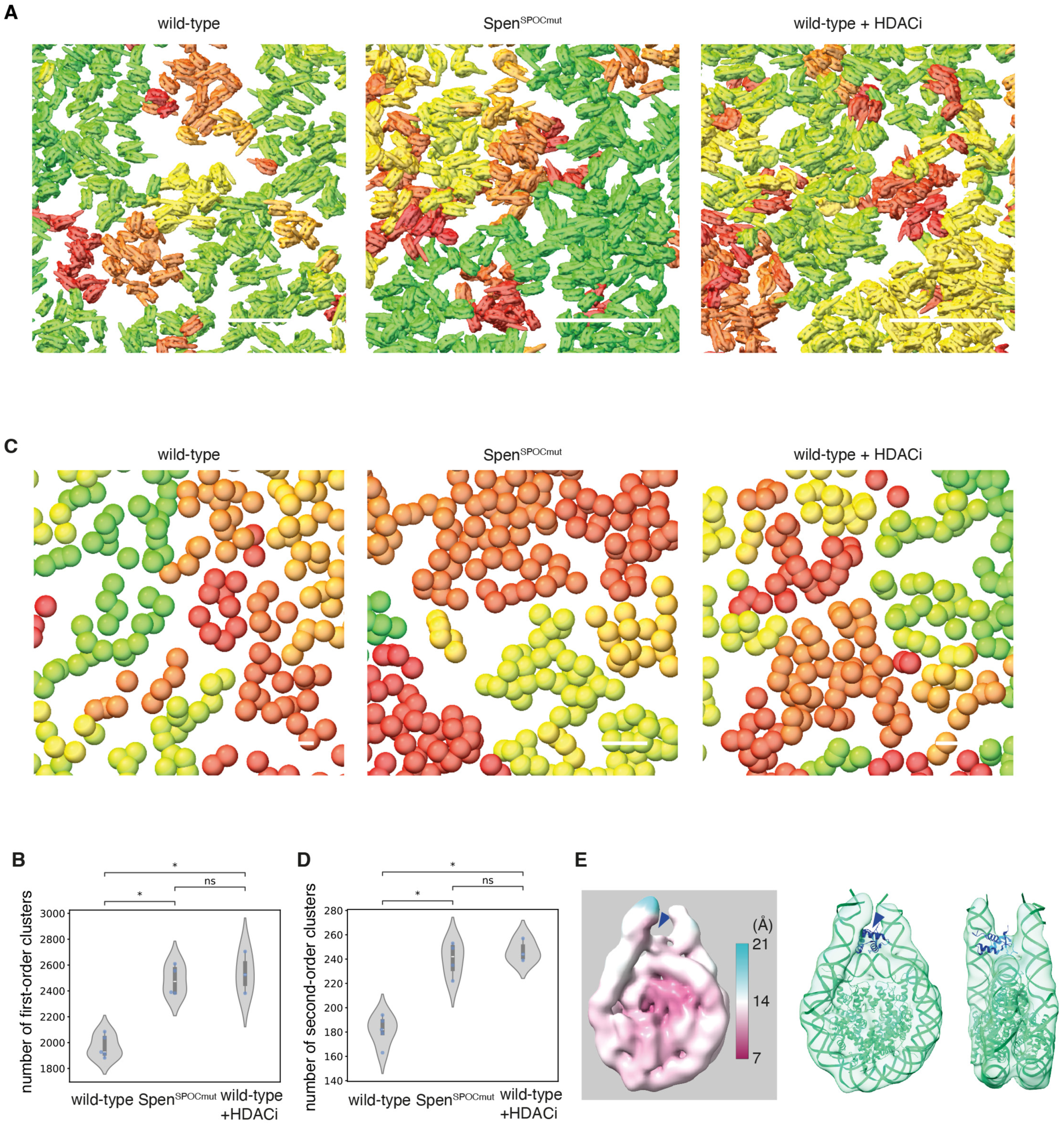
**(A)** Representative cropped views showing nucleosome-clutch clustering of day-7 differentiated wild-type, Spen^SPOCmut^, and HDACi-treated wild-type Xi within the tomogram volume (z = 46.6 nm) using H-DBSCAN. Nucleosomes belonging to the same cluster are depicted in the same colour shade. Scale bar = 50 nm. **(B)** A violin plot of the number of first-order clusters within the tomogram volume. **(C)** Representative cropped views showing chromatin-domain clustering of day-7 differentiated wild-type, Spen^SPOCmut^, and HDACi-treated wild-type Xi within the tomogram volume (z = 46.6 nm) using H-DBSCAN. Each nucleosome clutch (first-order cluster) is depicted as a sphere with a diameter of 28 nm around the centroid of the cluster, for simplification. Nucleosome clutches belonging to the same chromatin domain are depicted in the same colour. Scale bar = 50 nm. **(D)** A violin plot of the number of second-order clusters within the tomogram volume. **(E)** Left: STA map of the chromatosome of the Spen^SPOCmut^ Xi, coloured by local resolution. Extra density corresponding to histone H1 is indicated by arrows. Right: the human chromatosome model (PDB: 7DBP) was fitted into our STA map. Extra cryo-EM density corresponding to histone H1 (shown in navy in the fitted model) is indicated by arrows. **(B)** and **(D)**: each dot represents a data point from a subvolume (839 × 839 × 46.6 nm^3^) of an independent tomogram, consisting of >20,000 extracted nucleosome particles. Exact P-values between all data points are provided in Supplementary Data Table 1. N = 5 (wild-type); 5 (Spen^SPOCmut^); 3 (wild-type + HDACi).

**Extended Data Table 1.** Cryo-ET data acquisition parameters and STA map information.

| **Map type** | **wild-type,**<br>**Xi, day5**<br>**(EMD-59330)** | **wild-type,**<br>**Xi, day7**<br>**(EMD-59329)** | **wild-type,**<br>**other, day7**<br>**(EMD-59331)** | **Spen<sup>SPOCmut</sup>,**<br>**Xi, day7**<br>**(EMD-59292)** |
| --- | --- | --- | --- | --- |
| data acquisition |  |  |  |  |
| **Microscope** | Titan Krios G4 | Titan Krios G4 | Titan Krios G4 | Titan Krios G4 |
| **Voltage (kV)** | 300 | 300 | 300 | 300 |
| **Camera** | Falcon 4i | Falcon 4i | Falcon 4i | Falcon 4i |
| **Magnification** | 53000 | 53000 | 53000 | 53000 |
| **calibrated pixel**<br>**size (Å/px)** | 2.33 | 2.33 | 2.33 | 2.33 |
| **targeted total**<br>**electron dose**<br>**(e-/Å<sup>2</sup>)** | 125 | 150 | 150 | 125 |
| **targeted defocus**<br>**range (µm)** | -2 to 4 | -2 to 4 | -2 to 4 | -2 to 4 |
| **Automation**<br>**software** | Tomo5 | Tomo5 | Tomo5 | Tomo5 |
| STA analysis |  |  |  |  |
| **# of tomograms** | 3 | 6 | 5 | 4 |
| **Initial # of**<br>**nucleosomes** | 52195 | 108045 | 121400 | 125325 |
| **final # of particles** | 13584 | 19631 | 16780 | 14262 |
| **Map resolution (Å)**<br>**(FSC 0.143)** | 11.13 | 12.14 | 12.14 | 11.78 |

## Supplementary information

### Supplementary Methods

#### ChIP-seq for H3K27ac

Native ChIP-seq was performed largely as described in our previous work^10^ with all buffers supplemented with 5mM of the deacetylase inhibitor sodium butyrate (Sigma). 5×10^7^ mESCs were used per experiment. Briefly, cells were lysed in RSB (10mM Tris pH8, 10mM NaCl, 3mM MgCl_2_, 0.1% NP40) for 5 minutes on ice with gentle inversion before nuclei collection by centrifugation (1500g for 5 minutes at 4°C). Nuclei were resuspended in 1ml of RSB + 0.25M sucrose + 3mM CaCl_2_, treated with 200U of MNase (Fermentas) for 5 minutes at 37°C, quenched with 4µl of 1M EDTA, then centrifuged at 2000g for 5 minutes. The supernatant was transferred to a fresh tube as fraction S1. The remaining chromatin pellet was incubated for 1 hour in 300µl of nucleosome release buffer (10mM Tris pH7.5, 10mM NaCl, 0.2mM EDTA), carefully passed five times through a 27G needle, and then centrifuged at 2000g for 5 minutes. The supernatant from this S2 fraction was combined with S1 to make the final soluble chromatin extract. For each ChIP reaction, 100µl of chromatin was diluted in Native ChIP incubation buffer (10mM Tris pH 7.5, 70mM NaCl, 2mM MgCl_2_, 2mM EDTA, 0.1% Triton) to 1ml and incubated with anti-H3K27ac antibody (rabbit polyclonal, Abcam ab4729) overnight at 4°C.

The following day, samples were incubated for 1 hour with 40µl protein A agarose beads pre-blocked in Native ChIP incubation buffer with 1mg/ml BSA and 1mg/ml yeast tRNA, then washed a total of four times with Native ChIP wash buffer (20mM Tris pH7.5, 2mM EDTA, 125mM NaCl, 0.1% Triton-X100) and once with TE pH7.5. All washes were performed at 4°C. The DNA was eluted from beads by resuspension in elution buffer (1% SDS, 100mM NaHCO_3_) and shaking at 1000rpm for 30 minutes at 25°C, and was purified using the ChIP DNA Clean and Concentrator kit (Zymo Research). ChIP enrichment of H3K27ac was confirmed by qPCR comparing the *Nanog* promoter (marked by active chromatin modifications) to a gene desert region and SensiMix SYBR (Bioline, UK). 25-100ng of ChIP DNA was used for next-generation sequencing (NGS) library prep using the NEBNext Ultra II DNA Library Prep Kit with NEBNext Single indices (E7645). NGS libraries were quantified using by Qubit fluorometer (Invitrogen) and DNA fragment sizes of ∼200-800bp verified with Bioanalyzer 2100 (Agilent) with High Sensitivity DNA chips. Additional rounds of clean-up and/or size selection were performed if necessary, using Agencourt AMPure XP beads (Beckman Coulter) to remove residual adaptors or large (>1000bp) fragments. 2×81 paired-end sequencing was performed on an NextSeq500 (FC-404-2002) machine using NextSeq 500/550 High-Output v2.5 kits (150 cycles) (Illumina).

#### ChIP qPCR primers

*Nanog* promoter:

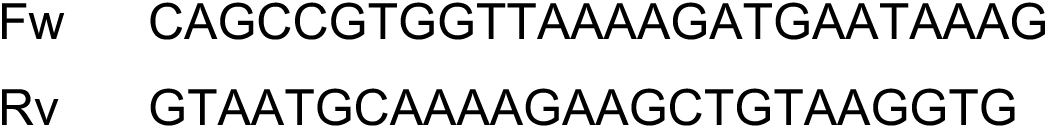

Gene desert:

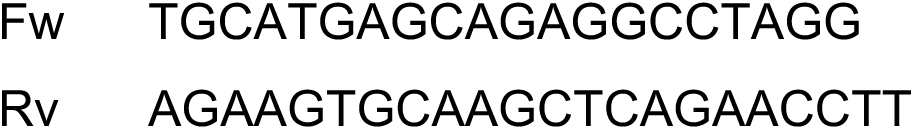

#### Voronoi density calculation

The logic of using Voronoi density calculation was adapted from previous studies^18,19^, and a custom Python script^68^ was developed to apply the analysis to nucleosome particle lists with assistance from ChatGPT 5.5 (OpenAI). Three-dimensional particle coordinates were read from STAR files and converted from pixels to nanometres using the supplied pixel size. A centred cuboidal region of interest was selected for each tomogram, and tomograms containing fewer than 20 particles after cropping were excluded. A three-dimensional Voronoi tessellation was calculated from the retained coordinates. The volume of each bounded Voronoi cell was determined from the convex hull of its vertices; unbounded cells and cells for which a valid positive volume could not be calculated were excluded. Voronoi-based local density was defined as the inverse of Voronoi-cell volume and expressed in nm. For each tomogram, the empirical cumulative distribution function of local density was evaluated on a common density axis spanning the density range across all included tomograms and experimental groups. Group-mean CDFs were calculated pointwise across tomograms, and the shaded regions represent the standard error of the mean.

**Supplementary Video 1: A tomogram of Xi, day-7 differentiated mESC**

Scale bar = 50 nm

**Supplementary Video 2: A tomogram of Xi, day-5 differentiated mESC**

Scale bar = 50 nm

**Supplementary Video 3: A tomogram of other chromosome, day-5 differentiated mESC**

Scale bar = 50 nm

**Supplementary Video 4: A tomogram of other chromosome, day-7 differentiated mESC**

Scale bar = 50 nm

**Supplementary Video 5: A tomogram of Xi, day-7 differentiated Spen^SPOCmut^ mESC**

Scale bar = 50 nm

**Supplementary Video 6: A tomogram of Xi, day-7 Differentiated, HDACi-treated wild-type mESC**

Scale bar = 50 nm

**Supplementary Table 1: P-values of plotted data**

## Notes

### Competing Interest Statement

The authors have declared no competing interest.

## References

1 Vinayak, V., Lakadamyali, M. & Shenoy, V. B. Architecture and regulation of nanoscale chromatin domains. Nat Commun, doi:10.1038/s41467-026-71213-5 (2026).

2 Liu, J., Ali, M. & Zhou, Q. Establishment and evolution of heterochromatin. Ann N Y Acad Sci 1476, 59–77, doi:10.1111/nyas.14303 (2020).

3 Giorgetti, L. et al. Structural organization of the inactive X chromosome in the mouse. Nature 535, 575–579, doi:10.1038/nature18589 (2016).

4 Deng, X. et al. Bipartite structure of the inactive mouse X chromosome. Genome Biol 16, 152, doi:10.1186/s13059-015-0728-8 (2015).

5 Loda, A., Collombet, S. & Heard, E. Gene regulation in time and space during X-chromosome inactivation. Nat Rev Mol Cell Biol 23, 231–249, doi:10.1038/s41580-021-00438-7 (2022).

6 Maeshima, K., Iida, S., Shimazoe, M. A., Tamura, S. & Ide, S. Is euchromatin really open in the cell? Trends Cell Biol 34, 7–17, doi:10.1016/j.tcb.2023.05.007 (2024).

7 Chen, J. K. et al. Nanoscale analysis of human G1 and metaphase chromatin in situ. EMBO J 44, 2658–2694, doi:10.1038/s44318-025-00407-2 (2025).

8 Tan, Z. Y. et al. Heterogeneous non-canonical nucleosomes predominate in yeast cells in situ. Elife 12, doi:10.7554/eLife.87672 (2023).

9 Hou, Z., Nightingale, F., Zhu, Y., MacGregor-Chatwin, C. & Zhang, P. Structure of native chromatin fibres revealed by Cryo-ET in situ. Nat Commun 14, 6324, doi:10.1038/s41467-023-42072-1 (2023).

10 Nesterova, T. B. et al. Systematic allelic analysis defines the interplay of key pathways in X chromosome inactivation. Nat Commun 10, 3129, doi:10.1038/s41467-019-11171-3 (2019).

11 Constantinescu, F. et al. Selective interaction of SMCHD1 with chromatin is governed by LRIF1 and SMCHD1 ATPase activity. Nat Commun, doi:10.1038/s41467-026-74427-9 (2026).

12 Jentink, N., Purnell, C., Kable, B., Swulius, M. T. & Grigoryev, S. A. Cryoelectron tomography reveals the multiplex anatomy of condensed native chromatin and its unfolding by histone citrullination. Mol Cell 83, 3236–3252 e3237, doi:10.1016/j.molcel.2023.08.017 (2023).

13 Gelleri, M. et al. True-to-scale DNA-density maps correlate with major accessibility differences between active and inactive chromatin. Cell Rep 42, 112567, doi:10.1016/j.celrep.2023.112567 (2023).

14 Gendrel, A. V. et al. Smchd1-dependent and -independent pathways determine developmental dynamics of CpG island methylation on the inactive X chromosome. Dev Cell 23, 265–279, doi:10.1016/j.devcel.2012.06.011 (2012).

15 Bowness, J. S. et al. Xist-mediated silencing requires additive functions of SPEN and Polycomb together with differentiation-dependent recruitment of SmcHD1. Cell Rep 39, 110830, doi:10.1016/j.celrep.2022.110830 (2022).

16 Teller, K. et al. A top-down analysis of Xa- and Xi-territories reveals differences of higher order structure at >/= 20 Mb genomic length scales. Nucleus 2, 465–477, doi:10.4161/nucl.2.5.17862 (2011).

17 Carignano, M. A. et al. Local volume concentration, packing domains, and scaling properties of chromatin. Elife 13, doi:10.7554/eLife.97604 (2024).

18 Otterstrom, J. et al. Super-resolution microscopy reveals how histone tail acetylation affects DNA compaction within nucleosomes in vivo. Nucleic Acids Res 47, 8470–8484, doi:10.1093/nar/gkz593 (2019).

19 Martinez-Sanchez, A., Baumeister, W. & Lucic, V. Statistical spatial analysis for cryo-electron tomography. Comput Methods Programs Biomed 218, 106693, doi:10.1016/j.cmpb.2022.106693 (2022).

20 Campello, R. J. G. B., Moulavi, D. & Sander, J. Density-Based Clustering Based on Hierarchical Density Estimates. 160–172 (2013).

21 Ricci, M. A., Manzo, C., Garcia-Parajo, M. F., Lakadamyali, M. & Cosma, M. P. Chromatin fibers are formed by heterogeneous groups of nucleosomes in vivo. Cell 160, 1145–1158, doi:10.1016/j.cell.2015.01.054 (2015).

22 Belyaev, N., Keohane, A. M. & Turner, B. M. Differential underacetylation of histones H2A, H3 and H4 on the inactive X chromosome in human female cells. Hum Genet 97, 573–578, doi:10.1007/BF02281863 (1996).

23 Jeppesen, P. & Turner, B. M. The inactive X chromosome in female mammals is distinguished by a lack of histone H4 acetylation, a cytogenetic marker for gene expression. Cell 74, 281–289, doi:10.1016/0092-8674(93)90419-q (1993).

24 Keohane, A. M., O’Neill L, P., Belyaev, N. D., Lavender, J. S. & Turner, B. M. X-Inactivation and histone H4 acetylation in embryonic stem cells. Dev Biol 180, 618–630, doi:10.1006/dbio.1996.0333 (1996).

25 Guenther, M. G., Barak, O. & Lazar, M. A. The SMRT and N-CoR corepressors are activating cofactors for histone deacetylase 3. Mol Cell Biol 21, 6091–6101, doi:10.1128/MCB.21.18.6091-6101.2001 (2001).

26 McHugh, C. A. et al. The Xist lncRNA interacts directly with SHARP to silence transcription through HDAC3. Nature 521, 232–236, doi:10.1038/nature14443 (2015).

27 Zylicz, J. J. et al. The Implication of Early Chromatin Changes in X Chromosome Inactivation. Cell 176, 182–197 e123, doi:10.1016/j.cell.2018.11.041 (2019).

28 Ariyoshi, M. & Schwabe, J. W. A conserved structural motif reveals the essential transcriptional repression function of Spen proteins and their role in developmental signaling. Genes Dev 17, 1909–1920, doi:10.1101/gad.266203 (2003).

29 Harrison, S. J. et al. A focus on the preclinical development and clinical status of the histone deacetylase inhibitor, romidepsin (depsipeptide, Istodax((R))). Epigenomics 4, 571–589, doi:10.2217/epi.12.52 (2012).

30 Rahman, F. et al. Mapping the nuclear landscape with multiplexed super-resolution fluorescence microscopy. Nat Commun 16, 6042, doi:10.1038/s41467-025-61358-0 (2025).

31 Toth, K. F. et al. Trichostatin A-induced histone acetylation causes decondensation of interphase chromatin. J Cell Sci 117, 4277–4287, doi:10.1242/jcs.01293 (2004).

32 Nozaki, T. et al. Dynamic Organization of Chromatin Domains Revealed by Super-Resolution Live-Cell Imaging. Mol Cell 67, 282–293 e287, doi:10.1016/j.molcel.2017.06.018 (2017).

33 Xia, J., Zhao, J. Z., Strom, A. R. & Brangwynne, C. P. Chromatin heterogeneity modulates nuclear condensate dynamics and phase behavior. Nat Commun 16, 6406, doi:10.1038/s41467-025-60771-9 (2025).

34 McInnes, L., Healy, J., Saul, N. & Großberger, L. UMAP: Uniform Manifold Approximation and Projection. Journal of Open Source Software 3, doi:10.21105/joss.00861 (2018).

35 Fung, H. K. H. et al. Genetically encoded multimeric tags for subcellular protein localization in cryo-EM. Nat Methods, doi:10.1038/s41592-023-02053-0 (2023).

36 Wang, Q., Mercogliano, C. P. & Lowe, J. A ferritin-based label for cellular electron cryotomography. Structure 19, 147–154, doi:10.1016/j.str.2010.12.002 (2011).

37 Shimazoe, M. A. et al. Linker histone H1 functions as a liquid-like glue to organize chromatin in living human cells. Science Advances 12, doi:ARTN eaec9801 10.1126/sciadv.aec9801 (2026).

38 Zhao, J. et al. H2AK119ub1 differentially fine-tunes gene expression by modulating canonical PRC1- and H1-dependent chromatin compaction. Mol Cell 84, 1191–1205 e1197, doi:10.1016/j.molcel.2024.02.017 (2024).

39 Matthews, R. E. et al. CRAMP1 drives linker histone expression to enable Polycomb repression. Mol Cell 85, 2503–2516 e2508, doi:10.1016/j.molcel.2025.05.031 (2025).

40 Miron, E. et al. Chromatin arranges in chains of mesoscale domains with nanoscale functional topography independent of cohesin. Sci Adv 6, doi:10.1126/sciadv.aba8811 (2020).

41 Jansz, N. et al. Smchd1 Targeting to the Inactive X Is Dependent on the Xist-HnrnpK-PRC1 Pathway. Cell Rep 25, 1912–1923 e1919, doi:10.1016/j.celrep.2018.10.044 (2018).

42 Wang, C. Y., Jegu, T., Chu, H. P., Oh, H. J. & Lee, J. T. SMCHD1 Merges Chromosome Compartments and Assists Formation of Super-Structures on the Inactive X. Cell 174, 406–421 e425, doi:10.1016/j.cell.2018.05.007 (2018).

43 Li, H. et al. Mapping chromatin structure at base-pair resolution unveils a unified model of cis-regulatory element interactions. Cell, doi:10.1016/j.cell.2025.10.013 (2025).

44 Shogren-Knaak, M. et al. Histone H4-K16 acetylation controls chromatin structure and protein interactions. Science 311, 844–847, doi:10.1126/science.1124000 (2006).

45 Chen, Y. et al. A Spectrum of Free Energy Landscape Topologies Encodes Chromatin Polymorphism and phase separation. biorxiv, doi:10.64898/2026.06.19.733383 (2026).

46 Zhou, H. et al. Multiscale structure of chromatin condensates explains phase separation and material properties. Science 390, eadv6588, doi:10.1126/science.adv6588 (2025).

47 Zhou, H. et al. Multi-scale structure of chromatin condensates rationalizes phase separation and material properties. Biorxiv, doi:10.1101/2025.01.17.633609 (2025).

48 Russell, K. et al. Near-atomistic simulations reveal the molecular principles that control chromatin structure and phase separation. bioRxiv, doi:10.1101/2025.11.17.688899 (2025).

49 Gibson, B. A. et al. Organization of Chromatin by Intrinsic and Regulated Phase Separation. Cell 179, 470–484 e421, doi:10.1016/j.cell.2019.08.037 (2019).

50 Langmead, B. & Salzberg, S. L. Fast gapped-read alignment with Bowtie 2. Nat Methods 9, 357–359, doi:10.1038/nmeth.1923 (2012).

51 Krueger, F. & Andrews, S. R. SNPsplit: Allele-specific splitting of alignments between genomes with known SNP genotypes. F1000Res 5, 1479, doi:10.12688/f1000research.9037.2 (2016).

52 Feng, J., Liu, T., Qin, B., Zhang, Y. & Liu, X. S. Identifying ChIP-seq enrichment using MACS. Nat Protoc 7, 1728–1740, doi:10.1038/nprot.2012.101 (2012).

53 Liao, Y., Smyth, G. K. & Shi, W. featureCounts: an efficient general purpose program for assigning sequence reads to genomic features. Bioinformatics 30, 923–930, doi:10.1093/bioinformatics/btt656 (2014).

54 Hagen, W. J. H., Wan, W. & Briggs, J. A. G. Implementation of a cryo-electron tomography tilt-scheme optimized for high resolution subtomogram averaging. Journal of Structural Biology 197, 191–198, doi:10.1016/j.jsb.2016.06.007 (2017).

55 Tegunov, D. & Cramer, P. Real-time cryo-electron microscopy data preprocessing with Warp. Nat Methods 16, 1146–1152, doi:10.1038/s41592-019-0580-y (2019).

56 Zheng, S., et al. AreTomo: An integrated software package for automated marker-free, motion-corrected cryo-electron tomographic alignment and reconstruction. biorxiv, doi:10.1101/2022.02.15.480593 (2022).

57 Chaillet, M. L., van Loenhout, J., Leung, M. R., Burt, A. & Tegunov, D. MissAlignment Teaches Itself Better Cryo-ET. doi:10.64898/2026.04.29.721716 (2026).

58 Cruz-Leon, S. et al. High-confidence 3D template matching for cryo-electron tomography. Nat Commun 15, 3992, doi:10.1038/s41467-024-47839-8 (2024).

59 Turonova, B. GAPStop(TM) - GPU Accelerated Python-base Stopgap for Template Matching doi:10.5281/ZENODO.10822455 (2024).

60 Sofroniew, N., et al. napari: a multi-dimensional image viewer for Python. doi:10.5281/zenodo.3555620 (2026).

61 makubans, T. turonova/cryoCAT: TANGO release. doi:10.5281/ZENODO.17817056 (2025).

62 Ermel, U. H., Arghittu, S. M. & Frangakis, A. S. ArtiaX: An electron tomography toolbox for the interactive handling of sub-tomograms in UCSF ChimeraX. Protein Sci 31, e4472, doi:10.1002/pro.4472 (2022).

63 Meng, E. C. et al. UCSF ChimeraX: Tools for structure building and analysis. Protein Sci 32, e4792, doi:10.1002/pro.4792 (2023).

64 Burt, A. et al. An image processing pipeline for electron cryo-tomography in RELION-5. FEBS Open Bio, doi:10.1002/2211-5463.13873 (2024).

65 Liu, Y. T. et al. IsoNet2 determines cellular structures at submolecular resolution without averaging. bioRxiv, doi:10.64898/2025.12.09.693325 (2025).

66 Maurer, V. J., Siggel, M. & Kosinski, J. PyTME (Python Template Matching Engine): A fast, flexible, and multi-purpose template matching library for cryogenic electron microscopy data. SoftwareX 25, doi:10.1016/j.softx.2024.101636 (2024).

67 Bellos, D., Choi, J. rosalindfranklininstitute/cryoCAT-better-peak-picking: v0.8.2.. doi:10.5281/zenodo.21828296 (2026).

68 Bellos, D. C., J. rosalindfranklininstitute/Nucleosome_Spatial_Analysis_supplementary: v0.0.3 [Computer software]. Zenodo, doi:10.5281/zenodo.22014939 (2026).

69 Tegunov, D., Xue, L., Dienemann, C., Cramer, P. & Mahamid, J. Multi-particle cryo-EM refinement with M visualizes ribosome-antibiotic complex at 3.5 A in cells. Nat Methods 18, 186–193, doi:10.1038/s41592-020-01054-7 (2021).

70 Soloviev, A. G., Murugova, T. N., Islamov, A. H. & Kuklin, A. I. FITTER. The package for fitting a chosen theoretical multi-parameter function through a set of data points. Application to experimental data of the YuMO spectrometer. Journal of Physics: Conference Series 351, doi:10.1088/1742-6596/351/1/012027 (2012).

71 Mann, H. B., Whitney, D. R. On a Test of Whether one of Two Random Variables is Stochastically Larger than the Other. The Annals of Mathematical Statistics 18, 50–60 (1947).

72 Bellos, D., Choi, J. rosalindfranklininstitute/Nucleosome_Spatial_Analysis: v0.0.4. doi:10.5281/zenodo.22014934 (2026).

